# Cell and tissue morphology determine actin-dependent nuclear migration mechanisms in neuroepithelia

**DOI:** 10.1101/536698

**Authors:** Iskra Yanakieva, Anna Erzberger, Marija Matejčić, Carl D. Modes, Caren Norden

**Affiliations:** Max Planck Institute of Molecular Cell Biology and Genetics, Pfotenhauerstraße 108, 01307 Dresden, Germany; Max Planck Institute for the Physics of Complex Systems, Nöthnitzer Straße 38, 01187 Dresden, Germany; Max Planck Center for Systems Biology, Pfotenhauerstraße 108, 01307 Dresden, Germany

**Author notes:** Corresponding Author: Caren Norden (C.N.).

## Abstract

Correct nuclear position is crucial for cellular function and tissue development. Depending on cell context however, cytoskeletal elements responsible for nuclear positioning vary. While these cytoskeletal mechanisms have been intensely studied in single cells, how nuclear positioning is linked to tissue morphology is less clear. Here, we compare apical nuclear positioning in zebrafish neuroepithelia. We find that kinetics and actin-dependent mechanisms of nuclear positioning vary in tissues of different morphology. In straight neuroepithelia nuclear positioning is controlled by Rho-ROCK-dependent myosin contractility. In contrast, in basally constricted neuroepithelia a novel formin-dependent pushing mechanism is found for which we propose a proof-of-principle force generation theory.

Overall, our data suggests that correct nuclear positioning is ensured by the adaptability of the cytoskeleton to cell and tissue shape. This in turn leads to robust epithelial maturation across geometries. The conclusion that different nuclear positioning mechanisms are favoured in tissues of different morphology highlights the importance of developmental context for the execution of intracellular processes.

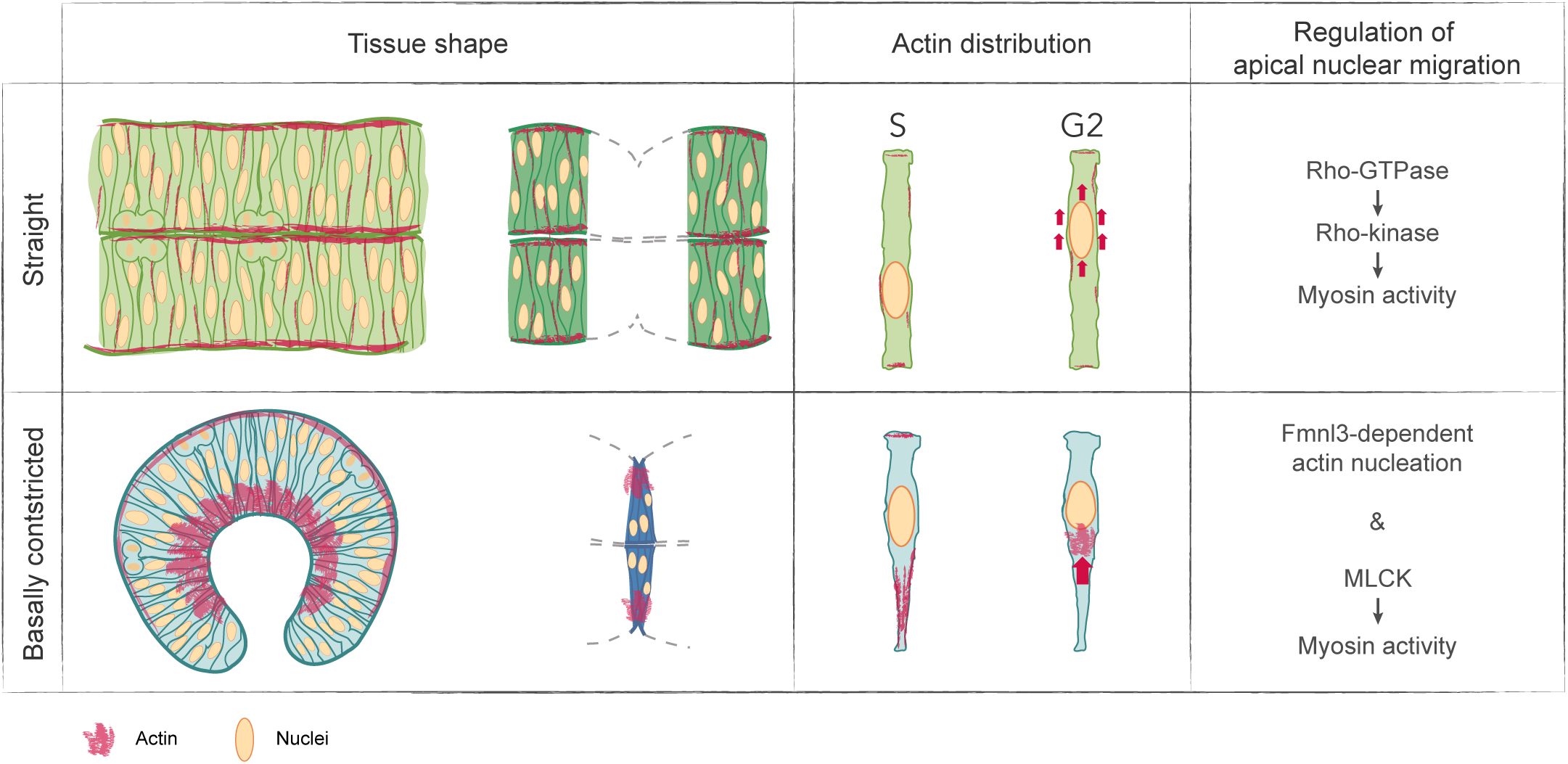

## INTRODUCTION

Nuclei can be positioned differently in cells depending on cell type, cell cycle phase, migratory state and differentiation stage (Gundersen and Worman, 2013). Nuclear positioning is a prerequisite for the correct execution of cellular programs including centered mitosis in fission yeast (Tran et al., 2001), differentiation of dermal cells in nematodes (Fridolfsson and Starr, 2010) or muscle cells in vertebrates (Roman and Gomes, 2018), and in neural system development (Shu et al., 2004; Tsai et al., 2007; Tsai and Gleeson, 2005). Due to its importance for correct cell function and tissue development, the position of the cell nucleus needs to be tightly controlled. To ensure exact positioning within cells, nuclei are actively transported by cytoskeletal elements and both actin (Gomes et al., 2005; Luxton et al., 2010) and microtubules (Fridolfsson and Starr, 2010; Reinsch and Gonczy, 1998; Tran et al., 2001) can exert pulling or pushing forces on nuclei using a variety of mechanisms. Interestingly, even within a single cell type, for example fibroblasts, the mechanisms of nuclear transport can differ depending on extracellular context (Levy and Holzbaur, 2008; Petrie et al., 2014; Wu et al., 2014). This striking variety of mechanisms not only underlines the importance of nuclear position regulation, but also illustrates the different means by which the cytoskeleton adapts to fulfill a precise task.

Diverse mechanisms of nuclear positioning have been studied extensively in cultured cells or the *C. elegans* zygote (Reinsch and Gonczy, 1998). However, how nuclear positioning is achieved in more complex settings such as tissues within developing organisms is not similarly well explored. In developing epithelia for example, complex shape changes occur at the single cell level and at the tissue scale. To date, it is not known how robust nuclear positioning is maintained across such varying cell- and tissue geometries.

Here, we address this question in pseudostratified neuroepithelia of the developing zebrafish. Pseudostratified neuroepithelia give rise to the nervous system, and correct nuclear positioning is crucial for their maturation. Nuclei in pseudostratified neuroepithelia are densely-packed and occupy different apicobasal positions in interphase (Lee and Norden, 2013; Sauer, 1935) when nuclear movements are stochastic (Kosodo et al., 2011; Leung et al., 2011; Norden et al., 2009). Preceding mitosis, however, nuclei are actively moved to the apical surface (Kosodo et al., 2011; Leung et al., 2011; Norden et al., 2009) (Fig. 1A). If nuclei fail to position apically, divisions occur at basal locations and these basally dividing cells perturb epithelial integrity and maturation (Strzyz et al., 2015). Interestingly, the cytoskeletal elements responsible for this crucial apical nuclear positioning differ depending on epithelium (Lee and Norden, 2013; Norden, 2017; Strzyz et al., 2016). In the extremely elongated cells of the rodent neocortex, movements are microtubule-dependent (Bertipaglia et al., 2017) and the mechanisms have been extensively studied (Hu et al., 2013; Shu et al., 2004; Tsai et al., 2010). In contrast, in shorter neuroepithelia nuclear positioning is driven by the actin cytoskeleton (Strzyz et al., 2016). However, the mechanisms by which actin generates the forces required for apical nuclear movement are still not fully understood. Rho-associated protein kinase (ROCK) has been implicated in apical nuclear migration (Meyer et al., 2011) in the *Drosophila* wing disc, but it is unclear whether this mechanism is conserved in other pseudostratified epithelia. Indications that nuclear positioning mechanisms might vary have come from a study of zebrafish retina and hindbrain neuroepithelia (Leung et al., 2011). However, how mechanisms differ and whether these differences are influenced by the tissue context remained elusive. Here, we investigate apical nuclear migration in zebrafish hindbrain and retinal neuroepithelia (Fig. 1B and B’). We reveal differences in nuclear kinetics between these tissues and show that these differences result from different actin-dependent mechanisms: in the hindbrain the Rho-ROCK pathway is involved in apical nuclear migration, while in the retina nuclear movements are driven by a formin-dependent pushing mechanism. We demonstrate that these mechanistic differences are conserved in other tissues, morphologically comparable to retina and hindbrain, and that migration modes can change when cell and tissue shape changes are induced.

**Figure 1.**
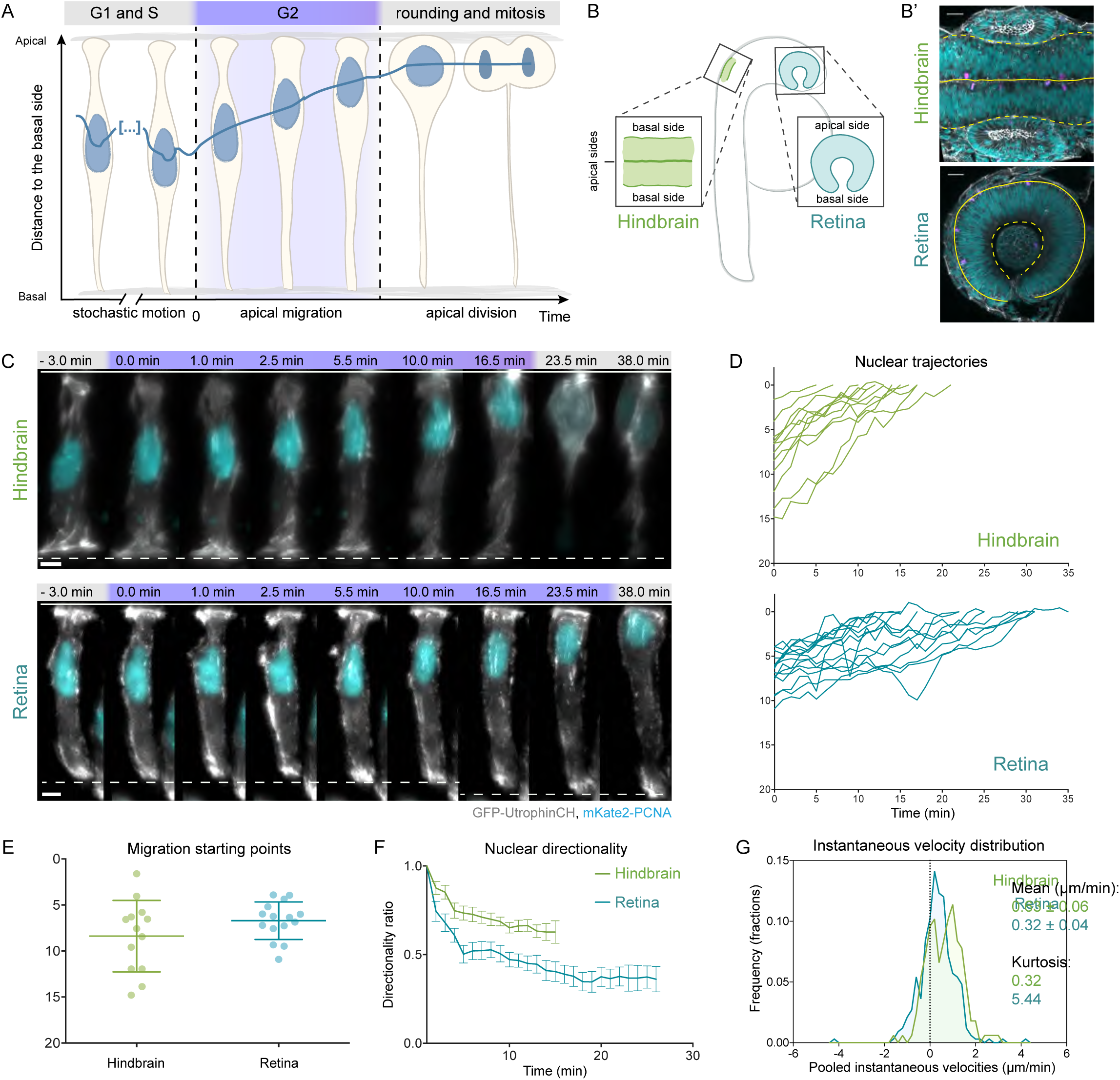
Apical nuclear migration in the hindbrain is faster and more directed than in the retina. (**A**) Neuroepithelial nuclei move stochastically in G1 and S and occupy diverse apico-basal positions. In G2 (highlighted in purple in the schematic and all montages), nuclei migrate to the apical side where cells divide. (**B**) Schematic of hindbrain and retinal neuroepithelia in the zebrafish embryo. Hindbrain is shown in green and retina in blue in all figures. (**B’**) Morphology of hindbrain at 18 hpf and retina at 24 hpf. Nuclear staining: DAPI (cyan), actin staining: phalloidin (gray), mitotic cells: pH3 (magenta). Solid lines mark the apical and dashed lines mark the basal tissue surface in all figures. (**C**) Example of apical nuclear migration in maximum projection of hindbrain and retinal cells, imaged with LSFM (Video 1). mKate2-PCNA labels nuclei (cyan), GFP-UtrophinCH labels actin (gray). (**D**) Apical migration trajectories. Start: 0 min = entry in G2. Finish: onset of cell rounding (nuclear position at cell rounding = 0 μm from the apical side). (**E**) Starting positions of hindbrain and retinal nuclei shown as mean ± SD. Variances comparison: p=0.0485, Levene’s test. (**F**) Directionality ratios shown as mean of all tracks, error bars represent SDs. Hindbrain = 0.63 ± 0.06, Retina = 0.36 ± 0.07. (**G**) Pooled instantaneous velocity distributions in hindbrain and retina. p<0.0001, Mann-Whitney test. Scale bars: (**B’**) 20 μm, (**C**) 5 μm.

## RESULTS

### Apical migration of retinal and hindbrain nuclei occurs with different kinetics

To understand the kinetic differences between apical nuclear migration in hindbrain and retinal neuroepithelia (Leung et al., 2011) (Fig. 1 B), we tracked nuclear movements in both tissues using light sheet fluorescent microcopy (LSFM) at sub-minute resolution (Icha et al., 2017) (Fig. 1 C). The G2-phase of the cell cycle, during which active nuclear migration occurs, was identified using a PCNA marker (Leung et al., 2011; Strzyz et al., 2015) (Fig. 1 C, Video 1). Nuclear trajectory analysis (Fig. 1 D) revealed that nuclei in the retina generally start G2 movements relatively close to the apical surface (Fig. 1 E) while the variance of starting points was greater in the hindbrain (Fig. 1 E, Table 1). Nevertheless, hindbrain nuclei migrated for shorter times than retinal nuclei (Table 1) indicating more directed movements. Quantitative analysis of apical nuclear movements confirmed that hindbrain nuclei indeed displayed a higher average instantaneous velocity compared to retinal nuclei (Table 1, Fig. 1 G and S4 H). Further, directionality ratio (Fig. 1 F, Table 1) and mean squared displacement analysis (Fig. S1 A) revealed a higher directionality of hindbrain nuclei (Table 1). Characterization of instantaneous velocity distributions showed that retinal nuclei more frequently undergo negative (basal) movements than hindbrain nuclei (Fig. 1 G). Comparing the kurtosis, a measure of the contribution of infrequent extreme deviations to the tails of the distribution, confirmed that extreme deviations were more common in retina than in hindbrain (Table 1, Fig. 1 G).

**Table 1.**
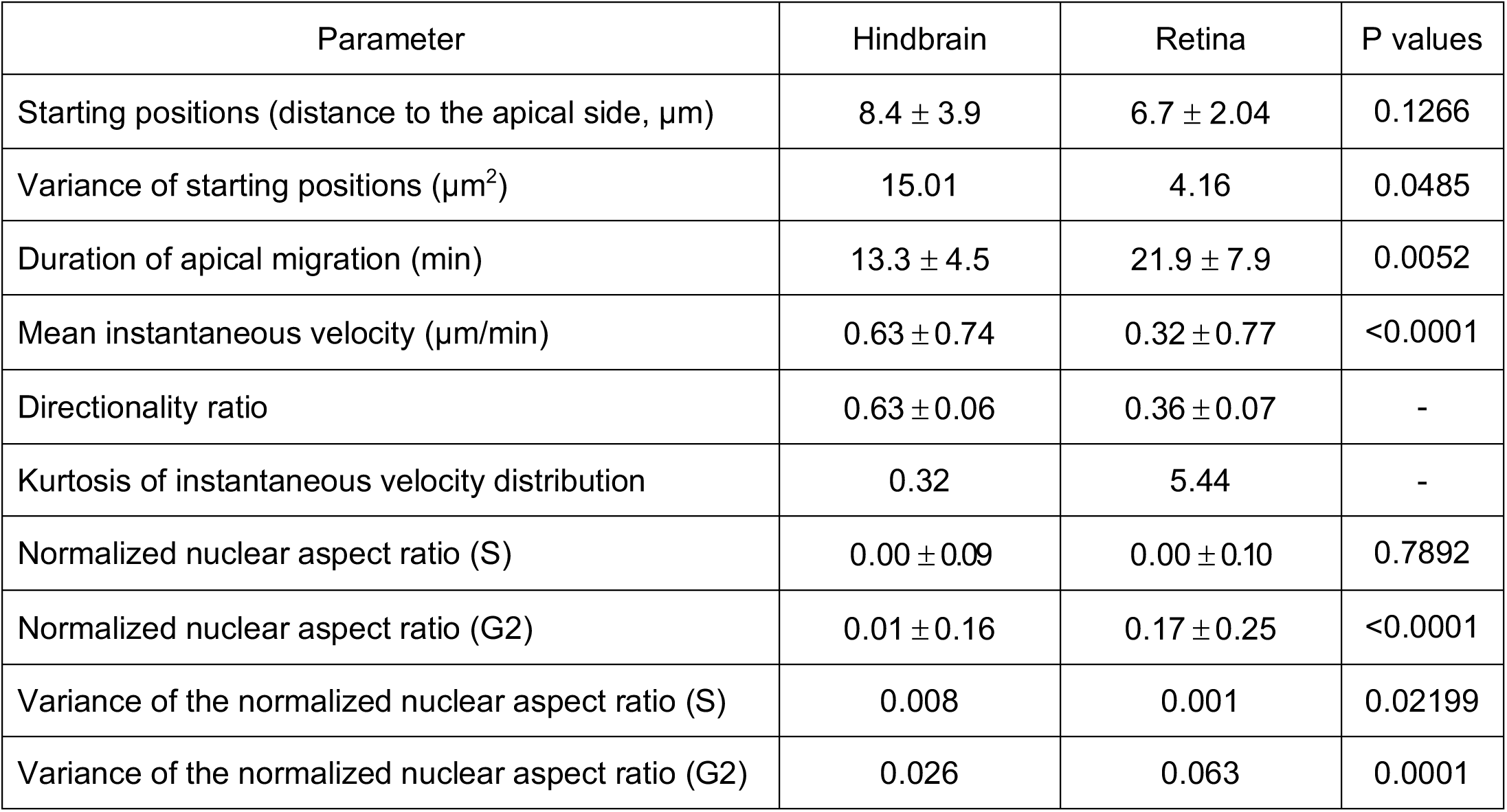
Nuclear migration kinetics and shape change parameters that differ between hindbrain and retina. Values shown represent mean ± SD. P values calculated using: Levene’s test for the variance of starting positions of migration and variances of the normalized nuclear aspect ratio; Mann-Whitney test for starting positions of migration, duration of apical migration, instantaneous velocity, and normalized nuclear aspect ratios.

Together, this data indicated that retinal nuclei move in a more saltatory manner than hindbrain nuclei.

Overall, our analysis confirms that apical nuclear movements differ between hindbrain and retina. While nuclei in the hindbrain start migrating from more variable apico-basal positions and move towards the apical surface in a directed and smooth manner, nuclei in the retinal neuroepithelium start more apically and their movements are slower and less directed.

These results made us speculate that the observed differences in the motion of hindbrain and retinal nuclei could result from different forces that move nuclei in the two tissues.

### Only retinal neuroepithelia nuclei show persistent aspect ratio changes during nuclear migration

If our hypothesis that different forces propel hindbrain and retinal nuclei during apical migration was correct, we assumed that we could infer these forces from nuclear deformations during movement. Nuclear deformability is inversely related to nuclear stiffness that depends predominantly on the expression of A-type lamins (Harada et al., 2014; Rowat et al., 2013; Swift et al., 2013). Different parts of developing mouse brain were shown to express low levels of Lamin A/C (Cho et al., 2019; Jung et al., 2012), which could suggest that in neuroepithelia nuclei are deformable and that their shape would change under the action of forces. To assess nuclear deformability in zebrafish neuroepithelia, we compared the expression levels of Lamin A/C and Lamin B1 in retinal and hindbrain nuclear envelopes. We found that while Lamin B1 was evenly distributed at nuclear envelopes of both tissues (Fig. 2 A), Lamin A/C was absent (Fig. 2 A) (Lamin A/C control staining (Fig. S2 B), suggesting that indeed neither hindbrain nor retinal nuclei were rigid. Live-imaging of nuclei expressing the nuclear envelope marker LAP2b corroborated this notion, revealing frequent deformations and dynamic indentations of nuclear envelopes (Fig. 2 B, Video 2). We analyzed these nuclear deformations for S- and G2-phase nuclei of both neuroepithelia by 3D nuclear segmentation (Fig. 2 C). In S-phase, nuclei in both tissues showed elongated, ellipsoidal shapes (Fig. 2 C). However, nuclei underwent frequent periods of deformation and relaxation fluctuating around the average with comparable variances (Fig. 2 D, S1 C, Table 1). While in G2 the aspect ratio of hindbrain nuclei fluctuated around a similar average as in S-phase, retinal nuclei persistently increased their aspect ratio in G2 and became more ovoid (Fig. 2 D, Table 1).

**Figure 2.**
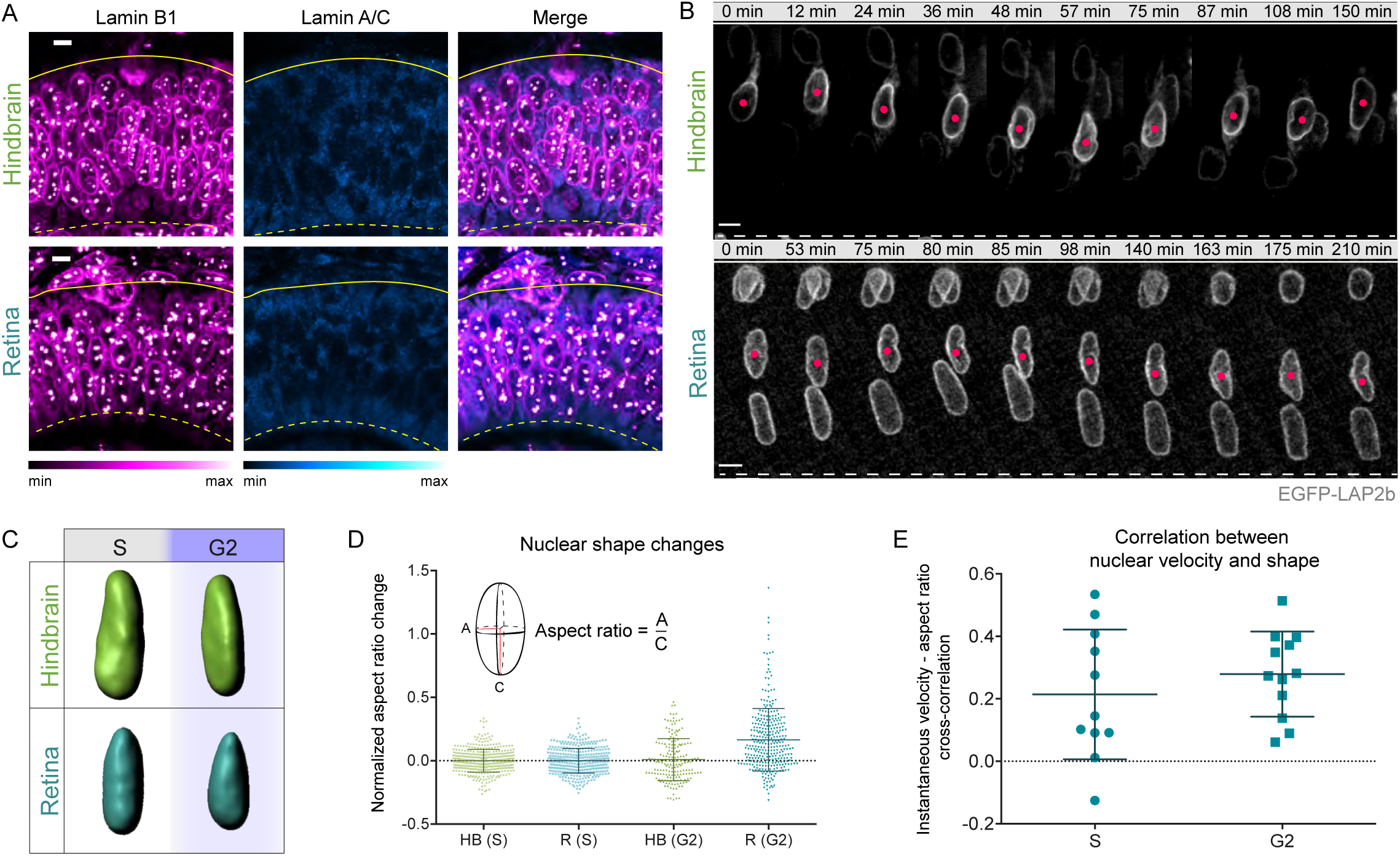
Hindbrain and retinal nuclei are deformable and experience different forces during apical nuclear migration. (**A**) Immunostaining of Lamin B1, and Lamin A/C in hindbrain and retinal cells (lookup tables indicate minimal and maximal signal values). (**B**) Dynamics of nuclear deformations in interphase hindbrain and retinal cells, visualized with nuclear envelope marker EGFP-LAP2b (Video 2). (**C**) 3D segmentation of hindbrain and retinal nuclei in S and G2. (**D**) Aspect ratio variability in S and G2 hindbrain and retinal cells. S-phase nuclei display similar variances of nuclear aspect ratio (p=0.7892, Levene’s test, Table 1). In G2, the variance and the mean aspect ratio increase more strongly for retinal nuclei (p<0.0001, Levene’s test and p<0.0001, Mann-Whitney test, respectively). (**E**) Cross-correlation analysis of nuclear instantaneous velocity and aspect ratio changes in the retina shows increased average correlation in G2. Error bars: SD. Scale bars: 5 μm.

Cross-correlation analysis showed that correlations between the dynamics of nuclear aspect ratio and instantaneous velocities in the retina were on average higher in G2 than in S-phase (Fig. 2 E, S1 D). The correlation between nuclear velocity and shape changes in retinal G2 nuclei suggested that both are caused by fluctuations in the force that propels retinal nuclei apically. One possible explanation for the shortening of retinal nuclei during migration was that they were being pushed to the apical side by a force originating basally of the nucleus. To test this notion, we performed laser ablation of a circular region in the center of hindbrain and retinal S and G2 nuclei (Fig. S1 E). To identify cell cycle phase, nuclei were labelled with GFP-PCNA (Fig. S1 F). H2B-RFP was used as a chromatin marker to visualize changes in the shape of the ablated regions (Fig. S1 F and G). In S-phase, the ablated regions of all retinal and hindbrain nuclei remained circular (Fig. S1 G). Similarly, no ablated region deformation was observed for G2 hindbrain nuclei. In retinal G2 nuclei, however, the ablated region frequently shortened in apico-basal direction and in some cases showed a basal indentation (Fig. S1 G and H) in agreement with a basal force acting on nuclei.

Together, this data indicated that nuclei in the retina are subjected to pushing forces in G2 while forces are evenly distributed during migration of G2 hindbrain nuclei.

### Distinct actomyosin pools are involved in nuclear movements in hindbrain and retina

To understand the different forces that drive apical nuclear migration in hindbrain and retina, we probed the cytoskeletal elements involved in their generation. In pseudostratified epithelia these forces have been shown to be generated by microtubules (Hu et al., 2013; Shu et al., 2004; Strzyz et al., 2016; Tsai et al., 2010) or actomyosin (Meyer et al., 2011; Norden et al., 2009). While it is known that in retinal neuroepithelia apical nuclear migration is actomyosin-dependent (Norden et al., 2009; Strzyz et al., 2015), the cytoskeletal elements that move nuclei in hindbrain neuroepithelia were yet unspecified. We thus performed Colcemid and Blebbistatin treatment to interfere respectively with microtubule or actomyosin activity. As shown before Blebbistatin but not Colcemid treatment impaired apical nuclear movement in the retina ((Norden et al., 2009; Strzyz et al., 2015) and Fig. S2 A, B, and C, Video 3). The same was true for the hindbrain (Fig. S2 A, B, and C, Video 3). This means that nuclear migration is microtubule-independent and actomyosin-dependent, in both neuroepithelia. The importance of the actin cytoskeleton in moving nuclei apically in both tissues was confirmed by combined interference with actin polymerization using Latrunculin A, and actin turnover, using Jasplakinolide (Fig. S2 D, Video 3).

We find that the same cytoskeletal elements drive nuclei to apical positions in retina and hindbrain, but their migration kinetics differ. We thus hypothesized that actin itself acts differently on nuclei of the two different epithelia leading to differences in migration kinetics and nuclear deformations. To test this idea, we investigated actin localization in hindbrain and retinal cells during S and G2. In hindbrain, the actin signal was mainly localized to the cell periphery in both cell cycle phases (Fig. 3 A, B, B’, S2 E, Video 4). In retinal tissue, actin was also seen in the cytoplasm of S-phase cells where the actin profile showed a high degree of variability (Fig. 3 A, S2 E, Video 4). During retinal G2 however, actin became considerably and more persistently enriched basally to the nucleus and the actin enrichment followed the organelle during migration (Fig. 3 A, B, B’, Video 4). Live imaging and analysis of myosin distribution also showed cytoplasmic enrichment basal to the nucleus in G2 retinal but not hindbrain cells (Fig. S2 F and G). Interestingly, the G2 actin ‘cloud’ following the nucleus in retinal cells did not have persistent intensity profiles but fluctuated with a similar frequency as the nuclear instantaneous velocity and the aspect ratio (Fig. 3 C, D, S2 H’, H’’, I, Video 4). Cross-correlation analysis showed a higher correlation of basal actin intensity with nuclear aspect ratio and lower with instantaneous velocity (Fig. 3 E). This suggested that basal actin enrichment is involved in the force-generation mechanism propelling retinal nuclei apically and that the same forces are responsible for the nuclear deformations that accompany retinal G2 movements. Together, these data demonstrate that actin and myosin are the cytoskeletal elements driving apical nuclear migration in both hindbrain and retina. However, different actin pools act during apical nuclear migration of hindbrain and retinal cells generating different forces that propel nuclei to the apical side.

**Figure 3.**
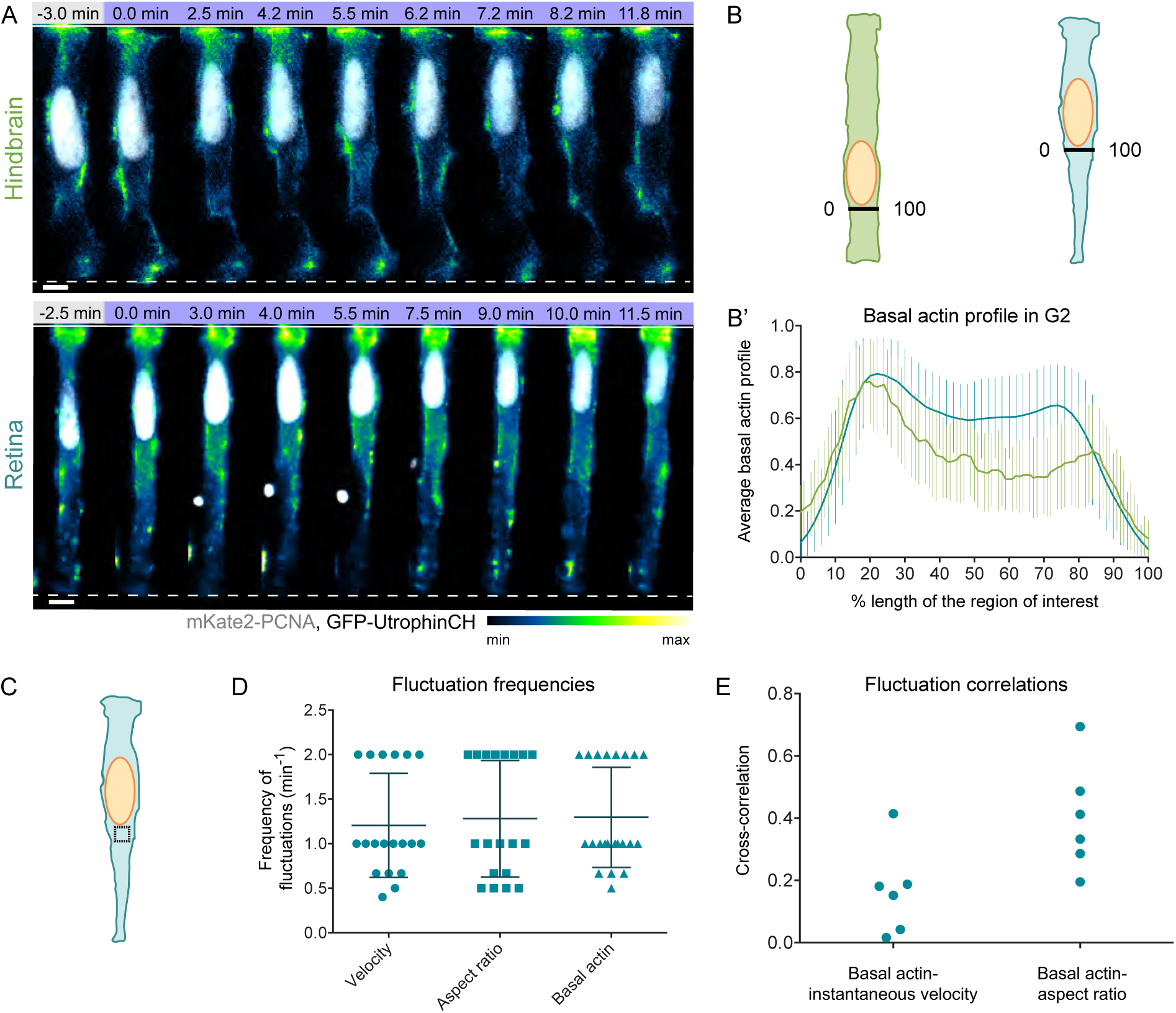
Distinct actomyosin pools are involved in apical nuclear migration in hindbrain and retina. (**A**) Actin distribution before and during apical migration in hindbrain and retinal cell (shown the central z plane of 3D stack) (Video 4). mKate2-PCNA labels nuclei (gray), GFP-UtrophinCH labels actin (lookup table indicates minimal and maximal GFP-UtrophinCH signal values). Scale bar: 5 μm. (**B**) Region basally of nucleus where GFP-UtrophinCH signal intensity was measured in the cells depicted in (A). (**B’**) Normalized average intensity distribution of GFP-UtrophinCH signal. Mean profile of all G2 time points is shown, error bars: SD. (**C**) Region where average GFP-UtrophinCH fluorescence intensity was measured basally to nucleus in a z-projection summing the intensities of all slices. (**D**) Pooled fluctuation frequencies of instantaneous velocity, nuclear aspect ratio, and basal actin intensity for the same retinal cell. Error bars: SD. (p=0.46 and p=0.05 for the pairs velocity-actin and aspect ratio-actin respectively, Mann-Whitney test). (**E**) Cross-correlation analysis between the fluctuations in basal actin and instantaneous velocity, as well as basal actin and nuclear aspect ratio.

### Different actomyosin regulators control apical nuclear movements in hindbrain and retinal cells

To dissect how actomyosin is controlled differently during apical nuclear migration in the different neuroepithelial cells, we investigated its upstream regulators. One pathway that regulates both myosin activity and actin polymerization, the Rho-GTPase – ROCK pathway was previously suggested to be involved in apical nuclear migration in the *Drosophila* wing disk (Meyer et al., 2011). We thus interfered with RhoA-GTPase or its effector ROCK, using respectively the small molecule inhibitors Rhosin, combined with Y16, (Fig. S3 A, Video 6) and Rockout (Fig. 4 B, Video 5). Compared to controls (Fig. 4 A and A’), both treatments led to impaired apical nuclear movement in hindbrain. Some G2 nuclei never reached the apical surface, resulting in basal mitosis (Fig. 4B, B’, D, S3A Video 5 and 6). Surprisingly, however, this was not the case in the retina. Here, nuclei moved like in controls (Fig. 4, B, B’, D, S3A, Video 5 and 6). The results were confirmed by overexpression of a dominant negative ROCK variant, DN-Rok2 (Marlow et al., 2002) (Fig. 4 F, Video 7). Hence, apical migration in the hindbrain but not in the retina depends on ROCK-dependent activation of myosin.

**Figure 4.**
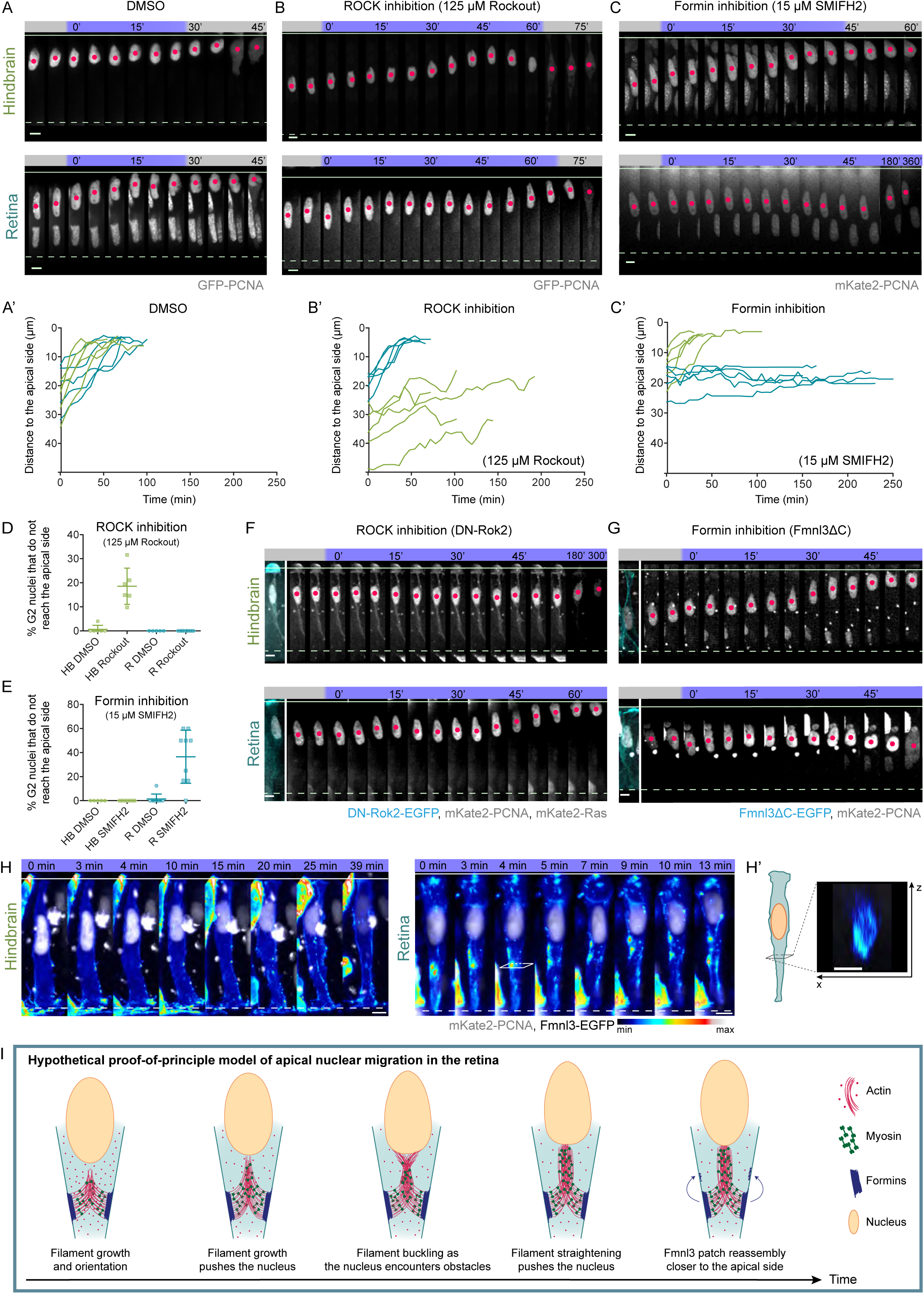
Different actomyosin regulators control apical nuclear movements in hindbrain and retinal cells. (**A**)-(**C**) Representative time-series of hindbrain and retinal cells treated with different actomyosin inhibitors (Video 5). (**A’**)-(**C’**) Representative trajectories of treated cells. Samples were incubated in (**A, A’**) DMSO, (**B**, **B’**) 125 μM Rockout (ROCK inhibitor), or (**C**, **C’**) 15 μM SMIFH2. (**D**), (**E**) Percentage of G2 nuclei in samples treated with (**D**) Rockout or (**E**) SMIFH2 unable to reach the apical side in hindbrain and retina. Error bars: SD. (p=0.0001 for Rockout-treated and p=0.0050 for SMIFH2-treated samples, Mann-Whitney test) (**F**), (**G**) Representative time-series of hindbrain and retinal cells expressing heat shock induced (**F**) DN-Rok2-EGFP (cyan), GFP-PCNA (gray) or (**G**) Fmnl3ΔC-EGFP (cyan), mKate2-PCNA (gray) (Video 8). (**H**) Representative time-series of hindbrain and retinal cells expressing Fmnl3-EGFP (maximum intensity projection, lookup table indicates minimal and maximal Fmnl3-EGFP signal values) and mKate2-PCNA (gray) (Video 8). (**H’**) Orthogonal (x-z) cross-section of the retinal cell’s basal process. Fmnl3-EGFP is seen enriched basally of the retinal nucleus. Scale bars: 5 μm. (**I**) Proof-of-principle theoretical model of a pushing mechanism that could drive apical nuclear migration in the retina. (for details, see Discussion and Supplemental material).

Despite its independence of the Rho-GTPase – ROCK pathway, general myosin inhibition by blebbistatin stalls apical nuclear migration also in the retina (Norden et al., 2009). We therefore tested whether another common myosin activator, MLCK (Myosin light-chain kinase), was involved in nuclear movements by interference with MLCK activity using the inhibitor ML-7 (Fig. S3 B, Video 6). This treatment had no effect on apical nuclear migration in the hindbrain but led to basally stalled G2 nuclei in the retina (Fig. S3 B). As earlier studies showed that MLCK can be involved in the formation and contractility of distinct pools of actin stress fibers (Totsukawa et al., 2000), we tested the role of different actin nucleators in the process. Using the small molecule CK-666 to inhibit the Arp2/3 complex, responsible for branched actin nucleation, had no effect on apical nuclear migration in retinal or hindbrain neuroepithelia (Fig. S3 C, Video 6). This result was confirmed by overexpression of a dominant negative variant of the Arp2/3 activator NWASP (Icha et al., 2016a) (Fig. S4 A and B, Video 7). Another major actin nucleator protein family are formins. Application of the pan-formin inhibitor SMIFH2 had no effect on nuclear migration in hindbrain (Fig. 4 C, C’, Video 5). In contrast, nuclei in retinal tissues treated with SMIFH2 often stalled basally for hours or did not reach the apical surface at all (Fig. 4 C, C’, E, Video 5).

As formins are a diverse protein family, we set out to identify the formins involved in nuclear migration in the retinal tissue. One suggestion came from a transcriptomics analysis (Sidhaye and Norden, GSE124779) that showed Fmnl3 (formin-like 3 protein) expression in the retina, which was confirmed by *in situ* hybridization (Fig. S4 C). Expression of GFP-tagged Fmnl3 revealed that the protein forms patches localized predominantly at the cell cortex in retinal progenitors (Fig. 4 H, H’, Video 8). During G2 these Fmnl3 patches were enriched 5 μm - 10 μm basally of the nucleus (Fig. 4 H, H’, Video 8) indicating that the observed actin ‘cloud’ basally to retinal nuclei (Fig. 3 A) was nucleated by Fmnl3. No basal Fmnl3 patches were observed in hindbrain cells (Fig. 4 H). Specific interference with Fmnl3 activity by overexpression of a dominant negative version, Fmnl3ΔC (Phng et al., 2015), demonstrated that indeed Fmnl3 perturbation affected apical nuclear migration in retinal but not hindbrain tissue (Fig. 4 G, Video 7).

These results show that while actin is the main driver of apical nuclear migration in both neuroepithelia, different actin regulators are at play (Fig. S3 D). In hindbrain, apical nuclear movement depends on Rho-GTPase-ROCK while in the retina MLCK-dependent contractility and formin-dependent actin polymerization are involved. To grasp how this novel formin-dependent mechanism could generate force to propel nuclei apically, we used our measured parameters combined with parameters taken from the literature to generate a proof-of-principle theoretical model for apically directed pushing. This working model recapitulates the observed saltatory movement of retinal nuclei and their average velocity, and offers a possible explanation for the dependence of the process on formin and myosin activity (Fig. 4 I and see discussion for model details).

### Different actin-dependent mechanisms of apical nuclear migration are linked to different cell and tissue shape

We showed that hindbrain and retinal neuroepithelial cells employ distinct actomyosin-dependent force generation mechanisms to move nuclei. This is surprising taking into consideration that these tissues display a similar pseudostratified architecture, and exist at similar developmental stages in the same organism. One possible explanation why nevertheless different mechanisms are used comes from a previous hypothesis that postulates that single-cell morphology as well as tissue-wide parameters like tissue shape or thickness could influence the cytoskeletal mechanisms generating the forces for apical nuclear migration (Strzyz et al., 2016). We thus tested whether cell and tissue morphology could influence the force generation mechanism observed in retinal versus hindbrain neuroepithelia. To this end, we analyzed tissue thickness, cell shape (Fig. S4 D, E, Table 2), and tissue-wide actomyosin distribution in both neuroepithelia (Fig. 5 A, A’, B). While tissue thickness was similar in retina and hindbrain (Fig. S4 D, Table 2), cell shape differed. Hindbrain cells were cylindrical with comparable apical and basal cell surface area (Fig. S4 E, Table 2), whereas retinal cells had a more conical shape with the cells’ apical surface areas greater than the basal surface areas (Fig. S4 E, Table 2). Cell shape differences were also reflected by different neuroepithelial geometry; a straight epithelium in hindbrain and a basally constricted epithelium in the retina (Fig. 1 B, B’). To understand the possible connections between tissue shape and force generation mechanisms, we further asked whether tissue-wide actin and myosin distribution could be different in the two neuroepithelia. We found that the tissue-wide distribution of actin, myosin and nuclei along the apicobasal axis differed: In hindbrain actin and myosin were evenly distributed along the lateral cell borders with peak intensities at apical and basal surfaces and nuclei dispersed all along the apicobasal axis (Fig. 5 A’ and B). In retinal tissue however, a basal bias of actin and myosin existed ((Matejcic et al., 2018; Sidhaye and Norden, 2017), Fig. 5 A’ and B) that likely limited the access of nuclei to the basal portion of the tissue and thus lead to the formation of a nuclear exclusion zone (Fig. 5 A’ and B, (Matejcic et al., 2018)). To test whether the observed differences in tissue architecture and distribution of the actomyosin cytoskeleton influence apical nuclear movements, we investigated the process in other straight or basally constricted zebrafish neuroepithelia at the midbrain-hindbrain boundary (MHB). The MHB is basally constricted at a point, termed MHBC (C for constriction (Gutzman et al., 2008)) while its neighboring regions (referred to as MHBS) are straight (Fig. 5 C and D). Like retina and hindbrain, MHBS and MHBC had comparable thickness (Fig. S4 D, Table 2). In concert with our findings in retina and hindbrain, actomyosin and nuclear distribution in the basally constricted MHBC matched findings in retinal tissue, while MHBS were comparable to hindbrain (Fig. 5 D, E, Table 2).

**Figure 5.**
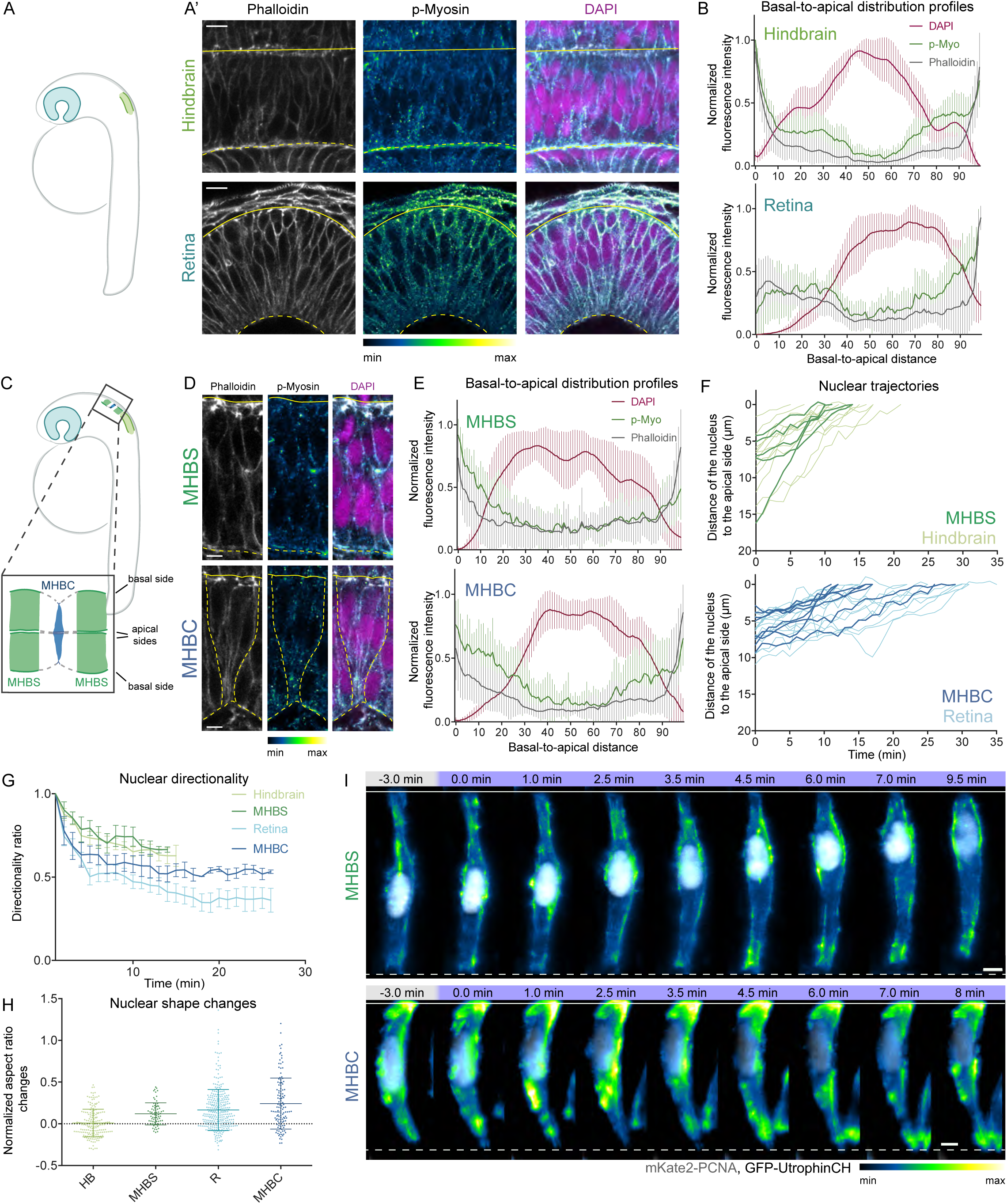
Different actin-dependent mechanisms of apical nuclear migration are linked to cell and tissue shape. (**A**) Schematic of the position and morphology of hindbrain and retinal neuroepithelia. (**A’**) Distribution of phalloidin (actin, gray), p-Myo (active (phosphorylated) myosin, lookup table indicates minimal and maximal signal values, and DAPI (nuclei, magenta) in hindbrain and retinal neuroepithelia. (**B**) Normalized average intensity distributions of phalloidin, p-Myo, and DAPI along the apicobasal axis of hindbrain and retinal neuroepithelium. The mean of all samples is shown, error bars: SD. (**C**) Schematic of the position and morphology of MHBS and MHBC neuroepithelia. MHBS: dark green, MHBC: dark blue. (**D**) Distribution of phalloidin (actin, gray), p-Myo (active (phosphorylated) myosin, lookup table indicates minimal and maximal signal values), and DAPI (nuclei, magenta) in MHBS and MHBC neuroepithelia. (**E**) Normalized average intensity distributions of phalloidin, p-Myo, and DAPI signal in MHBC and MHBS. The mean of all samples is shown, error bars: SD. (**F**) MHBS and MHBC nuclear trajectories compared to hindbrain and retinal trajectories. Hindbrain and retinal trajectories correspond to Fig. 1 D. (**G**) Directionality ratios of MHBS and MHBC nuclei. The mean of all tracks is shown with hindbrain and retinal data corresponding to Fig. 1 F. Error bars: SD. Final directionality ratios: MHBS = 0.67 ± 0.01, MHBC = 0.53 ± 0.01. (**H**) Average nuclear aspect ratio change in MHBS and MHBC with the onset of G2 (p=0.0221, Mann-Whitney test). Hindbrain and retinal aspect ratio changes correspond to Fig. 2 D. Error bars: SD. (**I**) Actin distribution before and during apical migration in MHBS and MHBC cells (shown maximum projection of 3D stack’s central 5 z-planes) (Video 9). mKate2-PCNA labels nuclei (gray), GFP-UtrophinCH labels actin (lookup table indicates minimal and maximal GFP-UtrophinCH signal values). Scale bars: (**A**) 10 μm, (**D**), (**I**) 5 μm.

**Table 2.** Tissue architecture, nuclear migration kinetics, and shape change parameters that differ between straight and curved neuroepithelial tissues. Values shown represent mean ± SD, all p values calculated using Mann-Whitney test, except for the variances of the normalized nuclear aspect ratio, for which Levene’s test was used.

| Parameter | Straight tissues |  | Curved tissues |  | P values |  |  |  |
| --- | --- | --- | --- | --- | --- | --- | --- | --- |
|  | Hindbrain | MHBS | Retina | MHBC | HB-MHBS | R-MHBC | HB-R | MHBS-MHBC |
| Cell length ($\mu\text{m}$) | $45.32 \pm 4.19$ | $38.27 \pm 4.6$ | $46.13 \pm 5.13$ | $41.54 \pm 4.6$ | 0.1017 | 0.0053 | 0.5780 | 0.0872 |
| Apical-to-basal footprint ratio | $1.25 \pm 0.47$ | $1.46 \pm 0.76$ | $2.32 \pm 0.76$ | $2.32 \pm 1.28$ | 0.1312 | 0.2617 | <0.0001 | <0.0001 |
| Duration of apical migration (min) | $13.3 \pm 4.5$ | $11.3 \pm 2.5$ | $21.9 \pm 7.9$ | $17.8 \pm 6.6$ | 0.3540 | 0.2457 | 0.0052 | 0.0326 |
| Mean instantaneous velocity ($\mu\text{m}/\text{min}$) | $0.63 \pm 0.74$ | $0.66 \pm 0.68$ | $0.32 \pm 0.77$ | $0.36 \pm 0.52$ | 0.9423 | 0.7831 | <0.0001 | 0.0038 |
| Directionality ratio | $0.63 \pm 0.06$ | $0.67 \pm 0.01$ | $0.36 \pm 0.07$ | $0.53 \pm 0.01$ | - | - | - | - |
| Normalized nuclear aspect ratio (G2) | $0.01 \pm 0.16$ | $0.12 \pm 0.13$ | $0.17 \pm 0.25$ | $0.24 \pm 0.31$ | <0.0001 | 0.0324 | 0.0001 | 0.0221 |
| Variance of the normalized nuclear aspect ratio (G2) | 0.026 | 0.017 | 0.063 | 0.096 | 0.0402 | 0.0007 | 0.0001 | <0.0001 |

Concluding that MHBS displayed similar cell and tissue morphology as hindbrain tissue while MHBC was similar to the retina, we tested whether nuclear migration characteristics reflected these architectural similarities. Indeed, nuclear trajectories in the two basally constricted neuroepithelia (MHBC and retina) showed striking resemblance, as did the trajectories in the two straight neuroepithelia (MHBS and hindbrain) (Fig. 5 F). The duration of apical nuclear migration in the straight tissues was shorter than in the two basally constricted tissues (Fig. S4 F, Table 2). Hence, G2 nuclei in MHBS moved faster than those in MHBC and mean instantaneous velocities between MHBS and hindbrain matched, as did velocities between MHBC and retina (Fig. S4 G, Table 2). Furthermore, nuclear movements in MHBS showed a higher directionality ratio than in MHBC (Fig. 5 G, Table 2) and MHBC nuclei, like retinal nuclei, changed their aspect ratio during G2 while the nuclear aspect ratio changed significantly less in MHBS cells (Fig. 5 H, Table 2). Further, live imaging of G2 actin showed a ‘cloud’ of actin accumulation following the nucleus in MHBC, similarly to actin in retinal cells (Fig. 5 I, Video 9) while in MHBS cells actin remained cortical during apical movement, similar to actin in hindbrain (Fig. 5 I, Video 9). To investigate whether also the pathways responsible for apical nuclear migration were conserved between tissues of similar morphology, we blocked Rho-Rock activity in MHBS and MHBC. We found that this treatment stalled apical nuclear migration only in MHBS, as seen for the morphologically similar hindbrain tissue but not in the MHBC (Fig. S4 H). This confirmed that tissue shape is linked to the mechanisms of apical nuclear migration.

We further tested whether a change in cell and tissue morphology would influence apical nuclear migration mechanisms. To this end, we used a previously published laminin *α*-1 (Pollard et al., 2006) morpholino that was shown to give rise to flat retinal neuroepithelia (Sidhaye and Norden, 2017). We confirmed that this treatment lead to changes in retinal cell shape in a dose-dependent manner (Fig. S4 E). Higher doses of the laminin *α*-1 morpholino resulted in straighter retinal tissue shape (Figure 6 A) and more cylindrical retinal cell shape (Fig. 6 B, S4E). In embryos treated with lower doses of laminin *α*-1 morpholino, a basal actin cloud was still observed in G2 and nuclei moved towards apical positions (n=4, Fig. 6 B). In embryos treated with higher morpholino dose in most cells no cytoplasmic actin enrichment was observed and actin was seen more laterally. In this case nuclei either did not move apically in G2 (n=2), or nevertheless reached apical positions, most likely using a different mechanism (n=3, Fig. 6 B).

**Figure 6.**
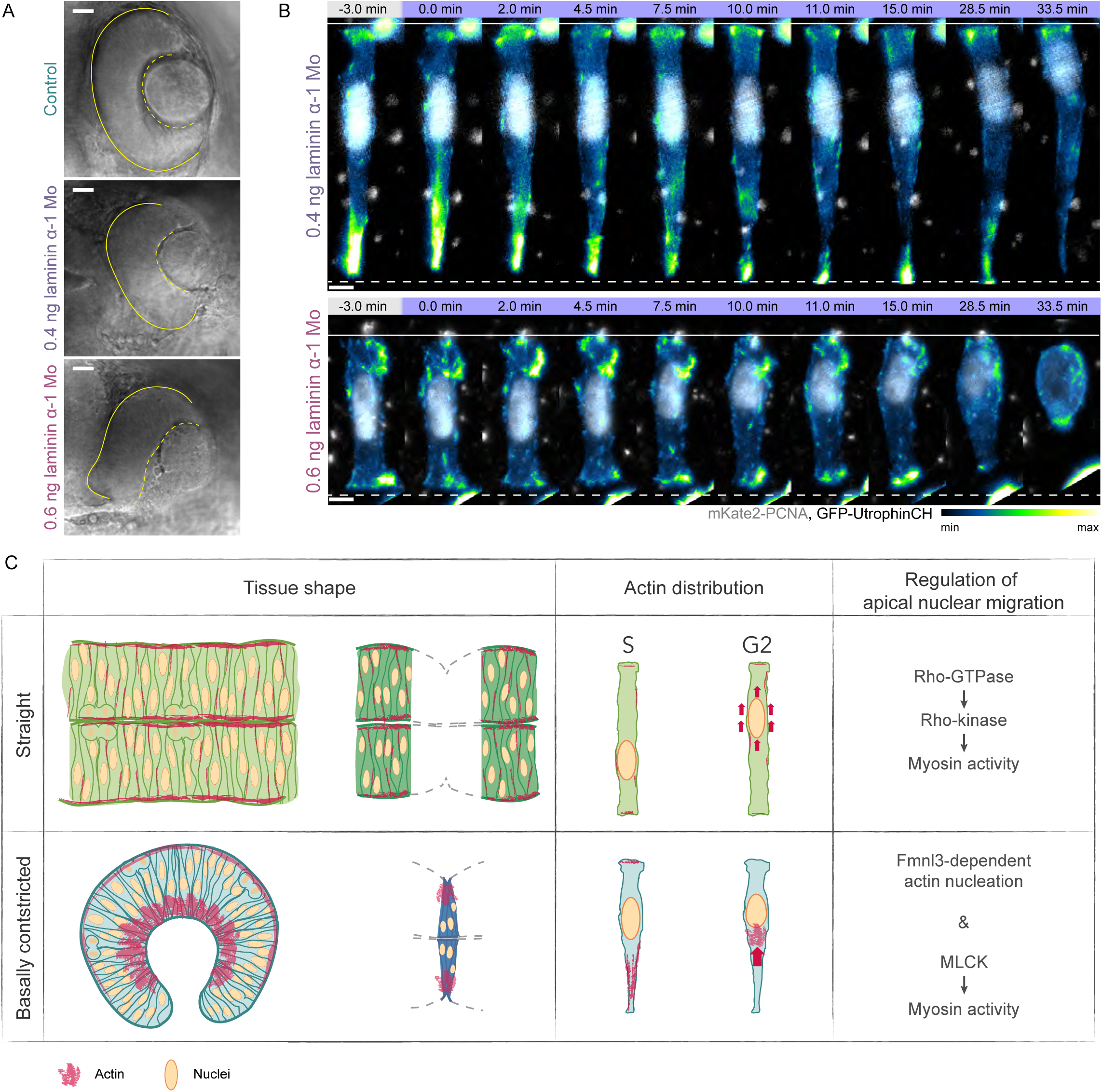
Tissue shape and the force generation mechanisms of apical nuclear migration are likely linked by the distinct actomyosin distribution in straight and basally constricted tissues. (**A**) Retinal tissue morphology in control and laminin α-1 morphant embryos, injected with different amounts of morpholino. (**B**) Representative time-series of retinal cell morphology and actin distribution during apical migration in laminin α-1 morphant embryos, injected with different amounts of morpholino (shown maximum projection of all z-planes in the 3D stack) (Video 9). mKate2-PCNA labels nuclei (gray), GFP-UtrophinCH labels actin (lookup table indicates minimal and maximal GFP-UtrophinCH signal values). Scale bars: (**A**) 10 μm, (**B**) 5 μm. (**C**) Schematic summary of suggested links between tissue shape and mechanisms of apical nuclear migration. Straight and basally constricted tissues show different distributions of actomyosin. In straight tissues actomyosin is evenly distributed along the lateral sides of cells. An enrichment of actomyosin is observed baso-laterally in basally constricted tissues. A basal actomyosin network that pushes the nucleus to the apical side could thus only be formed in cells of basally constricted, but not straight tissues.

These results indicate that indeed, cell and tissue shape are linked to apical nuclear migration mechanisms. When atypical shape changes occur within the tissue, the cytoskeletal machinery is able to at least partially adapt to ensure apical nuclear migration.

We conclude that cell and tissue shape influence the actin-dependent mechanisms that move nuclei apically in similar tissues within the same organism. The fact that actin is differently regulated in differently shaped neuroepithelia to achieve the same goal stresses the robustness of apical nuclear migration in diverse pseudostratified tissues.

## DISCUSSION

Our study demonstrates that the cytoskeletal force generating mechanisms for nuclear positioning vary in similar tissues within the same organism depending on cell and tissue geometry. In particular, we found that apical nuclear migration in pseudostratified neuroepithelia of different cell and tissue shape is driven by distinct actomyosin-dependent force generating mechanisms. We propose that an actomyosin contractility-dependent mechanism, downstream of the Rho-ROCK pathway, acts in the straight hindbrain tissue (Fig. 6 C). In contrast, in the retinal tissue that shows basally constricted morphology and a corresponding basal actomyosin accumulation, nuclei are possibly pushed by a formin-nucleated expanding actin network (Fig. 6 C). These mechanisms are conserved in tissues of similar shape and changes in cell morphology can lead to adaptation of mechanisms to ensure successful apical nuclear migration.

### Prerequisites and possible mechanisms of apical nuclear migration

We showed that nuclei in hindbrain and retina are highly deformable. This is most likely due to their lack of Lamin A/C expression, a major contributor to nuclear mechanical stiffness (Swift et al., 2013). Retinal and hindbrain nuclei display frequent deformations in all phases of the cell cycle further arguing that their nuclear envelopes are relatively soft. These deformations are most prominent during nuclear apical migration in the retinal neuroepithelium. Future studies are needed to test whether the observed deformability of nuclei in neuroepithelia enable the complex nuclear migration patterns occurring in these highly proliferative epithelia. With developmental progression, the epithelium becomes more and more crowded (Matejcic et al., 2018) which means that nuclei need to squeeze through increasingly confined spaces. Here, nuclear deformability could help to ensure successful apical migration.

Interestingly, the mechanisms driving apical nuclear migration vary depending on tissue: Apical nuclear migration in hindbrain cells is driven by ROCK activity. This could mean that nuclear movements depend on ROCK-induced cortical contractility as previously implicated in apical nuclear migration in *Drosophila* wing disc (Meyer et al., 2011). One possible contractility-dependent mechanism is the generation of cortical flows, similarly to those in *C. elegans* zygotes or cells undergoing adhesion-free migration (Bergert et al., 2015; Mayer et al., 2010; Munro et al., 2004). Gradual contraction of the cortex due to the action of a multitude of myosin motors could further explain the faster and smoother nuclear migration in hindbrain. Future experiments, including specific perturbation of cortical contractility, will help to test this hypothesis.

In the retina, basal enrichment of actin and myosin in G2 was reported previously (Leung et al., 2011; Norden et al., 2009) and it was proposed that nuclei are propelled by basal process constriction. However, we did not observe such constriction during apical migration of retinal nuclei (Fig. 1 C and 3 A, Video 1 and 4). In contrast, the periodic enrichment of actomyosin that closely follows nuclei during movement and the accompanying nuclear deformations argue that an expanding actin network pushes the nucleus apically.

Such pushing mechanism has not yet been described for nuclear migration in developing tissues. However, it has been shown that polymerizing actin networks can generate saltatory movements *in vitro* (Delatour et al., 2008), for the motility of intracellular parasites (Gerbal et al., 2000; Soo and Theriot, 2005) and during the pushing of chromosomes in mouse oocytes (Li et al., 2008; Yi et al., 2013).

To formalize how polymerizing actin could generate the forces necessary for nuclear migration in neuroepithelial cells, we developed a proof-of-principle theoretical model (Supplemental Material & Fig. 4 I):

Our mathematical model argues that cortex-anchored bundled f-actin could explain the observed phenomena. Cell geometry can direct f-actin growth, with frustum-shaped cells preferentially growing f-actin into the cytoplasm directed toward the larger side (Reymann et al., 2010). Thus, actin bundles growing via formin-catalyzed polymerization at the anchored end could push the nucleus toward the apical side by direct contact against the nuclear envelope. Intriguingly, the model shows that for appropriate parameter ranges the critical length for Euler buckling of the bundle (Bathe et al., 2008; Kierfeld et al., 2006) is consistent with the observed trailing distance of the formin enrichment zone behind the nucleus (Supplemental Material). Further, as the nucleus encounters spatial constraints, the effective dynamic viscosity would rise, resulting in a decreased buckling length threshold. As filaments then buckle, the continued addition of actin monomers at the trailing formin sites provide increased forces to the nucleus due to the rising stresses in the filaments, eventually squeezing the nucleus through. The buckled filaments and bundles then straighten, leading to a burst of increased velocity. A key prediction of this proof-of-principle model is that the average velocity of the nucleus is dominated by the speed of formin-catalyzed f-actin polymerization. Indeed, we observe that these are in close concert (Fig. 1 G and S4 G, Table 1, Supplemental Material). The possibility that the coordinated expansion of the actin network pushes the nucleus apically is an exciting new mechanism for positioning nuclei. It should be stressed however, that this model is currently based only partly on measured parameters and that some parameters, while consistent with the literature still need to be explored. Furthermore, at this point we explicitly do not exclude the possibility that force could also be generated by a contractility based mechanism. Further experiments that test and expand the current models will be important to clarify this point and will open exciting avenue for future investigations in neuroepithelia and beyond.

### The actin-dependent mechanisms employed for apical nuclear migration depend on tissue geometry

Our finding that neuroepithelial cells in tissues with different morphology use distinct actin-dependent mechanisms for apical nuclear movements demonstrates that the actomyosin cytoskeleton can use diverse means of force generation depending on cell and tissue context when performing a task critical for cell function and tissue development. Interestingly, actomyosin is responsible for both cell and tissue shape generation and maintenance, as well as the generation of the intracellular forces during nuclear positioning. Hindbrain and retinal neuroepithelia differ in cell and tissue shape, as well as tissue-wide cytoskeleton organization (Fig. 5 A, A’, B, (Matejcic et al., 2018; Sidhaye and Norden, 2017)). It is tempting to speculate that intracellular actin distribution, related to tissue shape, influences the force generation mechanism for apical nuclear migration. The tissue-wide basal enrichment of actomyosin, important for basal constriction and thereby development of the hemispheric retina (Nicolas-Perez et al., 2016; Sidhaye and Norden, 2017) could favor the formation of the basal structures that push the nucleus apically (Fig. 6 C). Such a mechanism would be efficient in tissues in which cells are constricted basally and subsequently show a basal actomyosin bias which is absent in the hindbrain, where a different mechanism is employed (Figure 6C). Of note, this basal actomyosin bias creates a region in the tissue that is inaccessible for nuclei and this could also explain the more apical starting points of nuclei in the retina compared to hindbrain (Matejcic et al., 2018).

To probe whether and how the cytoskeleton moves nuclei in other epithelial architecture regimes, more studies of apical nuclear migration are needed. It is for example not known whether mechanisms differ in apically constricted hindbrain regions or in tissues that thicken over development, like the more mature retinal neuroepithelium (Matejcic et al., 2018). Apical nuclear migration is a prerequisite for apical mitosis and, hence, correct tissue development in all pseudostratified epithelia. As these tissues are the precursors of organs in diverse organisms (Norden, 2017), cytoskeletal adaptability to move nuclei depending on the surroundings is most likely crucial for successful organogenesis. The variety of the described mechanisms of apical nuclear migration across systems suggests that, for pseudostratified epithelial cells, this important end justifies the many means.

## Materials and Methods

### Zebrafish husbandry

Wild type zebrafish were maintained and bred at 26°C. Embryos were raised at 21, 28.5, or 32°C in E3 medium. Animals were staged in hpf (hours post fertilization) according to (Kimmel et al., 1995). From 8 hpf the medium was supplemented with 0.2 mM 1-phenyl-2-thiourea (PTU) (Sigma-Aldrich) to prevent pigmentation. Embryos were anaesthetized in 0.04% tricaine methanesulfonate (MS-222; Sigma-Aldrich) prior to live imaging. Live imaging was performed for 6-14 hours from 18 hpf for the hindbrain and MHB, and from 24 hpf for the retina. All animal work was performed in accordance with EU directive 2010/63/EU, as well as the German Animal Welfare Act.

### RNA and DNA injections

To mosaically label cells in zebrafish neuroepithelia, DNA constructs were injected into one-cell stage embryos while mRNA was injected into the cytoplasm of a single blastomere of 32 - 128 cell stage embryos. DNA was injected at 10-25 pg per embryo. mRNA was synthesized using the Ambion mMessage mMachine kit and injected at 40 to 100 pg per embryo. The injection mix was prepared in water and the injected volume was 0.5-1.0 nl. A full list of the constructs used can be found in Table S1.

### Cloning strategies

Gateway cloning (Thermo Fisher Scientific) based on the Tol2 kit (Kwan et al., 2007) was used for all constructs.

#### pCS2+ mKate2-PCNA

Human PCNA coding sequence was amplified from pCS2+ GFP-PCNA (Leung et al., 2011) and a 3’-entry clone was generated. It was combined with mKate2 p5ENTR(L1-L2) (kind gift from Andrew Oates) and pCS2+ pDEST(R1-R3) (Villefranc et al., 2007).

#### T2-hsp70: EGFP-LAP2b

Rat LAP2b coding sequence was amplified from a pmRFP_LAP2beta_IRES_puro2b plasmid (Steigemann et al., 2009), Addgene, 21047) and a 3’-entry clone was generated. It was combined with hsp70 promoter p5ENTR(L4-R1) clone (Kwan et al., 2007), EGFP pMENTR(L1-L2) (Villefranc et al., 2007) and GW Tol2-pA2 p DEST backbone (Villefranc et al., 2007).

#### T2-hsp70: LMNA-mKate2

RNA was extracted from 24 hpf embryos using the TRI Reagent (Sigma-Aldrich) according to the manufacturers protocol. cDNA was synthesized using first strand cDNA synthesis kit (Fermentas/ Thermo-Fischer scientific). Zebrafish lamin A (BC163807.1) coding sequence was amplified from zebrafish cDNA to generate a middle entry clone without a stop codon at the end. The following primers were used:

5’ ggggacaagtttgtacaaaaaagcaggctggGAGTCGCAGCACACACTCTTT 3’

5’ ggggaccactttgtacaagaaagctgggtcAATAGAGCAGTTGTCCACTTTGG 3’ It was combined with hsp70 promoter p5ENTR(L4-R1) clone (Kwan et al., 2007), mKate2 p3ENTR(R2-L3) (kind gift from Andrew Oates) and GW Tol2-pA2 p DEST backbone (Villefranc et al., 2007).

#### T2-hsp70: Fmnl3ΔC-EGFP

The middle entry clone for truncated Fmnl3, lacking catalytic C terminus FH1, FH2, and DAD domains (pME-Fmnl3ΔC) (Phng et al., 2015) was a kind gift from Li-Kun Phng. It was combined with hsp70 promoter p5ENTR(L4-R1) clone (Kwan et al., 2007), EGFP p3ENTR(R2-L3) (Villefranc et al., 2007) and GW Tol2-pA2 p DEST backbone (Villefranc et al., 2007).

#### T2-hsp70: Fmnl3-EGFP

Zebrafish Fmnl3 (NM_001346154) coding sequence was amplified from zebrafish cDNA to generate a middle entry clone without a stop codon at the end. The following primers were used:

5’ ggggacaagtttgtacaaaaaagcaggctggATGGGGAATATTGAGAGTGTGG 3’

5’ ggggaccactttgtacaagaaagctgggtcGCAGATGGACTCGTCGAAGA 3’ It was combined with hsp70 promoter p5ENTR(L4-R1) clone (Kwan et al., 2007), mKate2 p3ENTR(R2-L3) (kind gift from Andrew Oates) and GW Tol2-pA2 p DEST backbone (Villefranc et al., 2007).

#### T2-hsp70: DN-Rok2-EGFP

The DN-Rok2 (Marlow et al., 2002) sequence was amplified to generate a middle entry clone. It was combined with hsp70 promoter p5ENTR(L4-R1) clone (Kwan et al., 2007), mKate2 p3ENTR(R2-L3) (kind gift from Andrew Oates) and GW Tol2-pA2 p DEST backbone (Villefranc et al., 2007).

### Heat shock of embryos

To induce expression of the heat shock promoter (hsp70) - driven constructs, embryos were incubated in a water bath at 17 hpf for imaging the hindbrain and at 23 hpf for imaging the retina to induce expression. The heat shock lasted 20 min at 37°C for Hsp70: DN-Rok2-EGFP and 30 min at 39°C for Hsp70: NWASP-CA-mKate2 and Hsp70:Fmnl3-EGFP. For induction of Hsp70:Fmnl3ΔC-EGFP, heat shocked lasted 15 minutes at 39°C for imaging the hindbrain neuroepithelium and for 20 minutes at 39°C when imaging the retinal neuroepithelium.

### Morpholino experiments

To knockdown gene function, the following amounts of morpholinos were injected into the yolk at one-cell stage:

0.4 or 0.6 ng laminin α-1 MO 5’TCATCCTCATCTCCATCATCGCTCA3’ (Pollard et al., 2006), 2 ng p53 MO 5’GCGCCATTGCTTTGCAAGAATTG3’ (Robu et al., 2007) to manage apoptosis.

### Drug treatments

All inhibitors were dissolved in DMSO, except Latrunculin A, that was dissolved in ethanol. Equal volumes of DMSO or ethanol as the stock inhibitor solution were used for control treatments. Dechorionated embryos were treated by incubation in E3 medium containing the inhibitors at their respective working concentrations (Table S2), either in plastic multi-well plates or in compartmentalized 35-mm glass bottom Petri dishes (Greiner Bio-One). All treatments were started after 17 hpf for the hindbrain and after 23 hpf for the retina.

#### Myosin perturbation for fixed imaging

Before fixation, embryos were incubated 6 h in DMSO, 125 μM Rockout, and 100 μM ML-7, and for 3 h in 100 μM Blebbistatin.

#### Live imaging of chemical perturbations

Embryos were dechorionated and pre-treated for one hour prior to mounting the sample. Concentrations for pre-treatment were 100 μM Rhosin, 25 μM Y16, 100 μM Rockout, 200 μM ML-7, 175 μM CK-666, and 10 μM SMIFH2. After mounting in agarose in glass-bottom dishes, embryos were incubated in inhibitor concentrations listed in (Table S2) for 14 hours and imaged using a spinning disk confocal microscope.

Cells that completed S-phase were counted using the CellCounter plugin in Fiji (Schindelin et al., 2012). Only embryos in which the drugs had an effect were analyzed. The numbers of treated and affected embryos are found in Table S3. Representative trajectories for controls, Rockout- and SMIFH2-treated cells were generated using the MTrackJ Fiji plugin.

### Immunofluorescence

For wholemount immunostainings embryos were fixed in 4% paraformaldehyde (PFA) (Sigma) in PBS for actomyosin and nuclear imaging, and Dent’s fixative for lamin immunostaining. The following antibodies and probes were used: Primary antibodies: 1:50 anti-phospho-myosin (Cell signaling), 1:500 anti-pH3 (Abcam), 1:200 anti-Lamin A/C (Abcam), 1:200 anti-Lamin B1 (Abcam). Secondary antibodies and fluorescent markers: 1:500 Alexa Fluor 647 anti-rat (Invitrogen), 1:500 Alexa Fluor 488 anti-Rabbit (Invitrogen), 1:500 Alexa Fluor 488 anti-mouse (Invitrogen), 1:500 Alexa Fluor 594 anti-Rabbit (Thermo Fischer Scientific), 1:50 Alexa Fluor 488 Phalloidin (Molecular Probes), 1:50 Rhodamine-Phalloidin (Molecular probes), DAPI.

### Microscope image acquisition

Experiments in the hindbrain were conducted between 18 hpf and 30 hpf, and between 24 hpf and 36 hpf in the retina before full onset of neurogenesis for each tissue.

#### Confocal scans

Fixed samples were imaged in Zeiss LSM 880 inverted point scanning confocal system (Carl Zeiss Microscopy) using the 40x/1.2 C-Apochromat water immersion objective (Carl Zeiss Microscopy). Samples were mounted in 1% agarose in glass bottom dishes (MatTek) or compartmentalized glass bottom dishes (Greiner Bio-One) filled with PBS and imaged at room temperature. z-stacks acquired had a thickness of 20-36 μm and step size of 0.75-1 μm. The microscope was operated by ZEN 2 (black edition) software.

#### Time-lapse imaging using spinning disk confocal microscope (SDCM)

Live imaging of apical migration perturbations was done using an Andor SDCM system. The spinning disk setup consisted of IX71 microscope (Olympus) and CSU-X1 scan head (Yokogawa). The samples were mounted in compartmentalized glass bottom dishes (Greiner Bio-One) or glass bottom dishes (MatTek) into 0.9% agarose in E3 medium containing 0.1 M HEPES pH=7.25 and 0.01% MS-222 (Sigma). The dish was filled with E3 medium containing 0.01% MS-222 (Sigma). Imaging was performed with UPLSAPO 60x/1.5 water immersion objective (Olympus) and Neo sCMOS camera (Andor) at 28.5°C regulated by an environmental chamber. A z stack of thickness 35-36 μm was acquired with 1 μm steps. Images were taken every 5 - 7 min for 12 - 14 hours. The microscope was operated by Andor iQ 3.0 software.

#### Time-lapse imaging using light-sheet fluorescent microscope (LSFM)

Imaging of single labelled cells in the hindbrain, retina, and MHB, was performed as previously described (Icha et al., 2016b) using the Lightsheet Z.1 (Carl Zeiss Microscopy). The sample chamber was filled with E3 medium containing 0.01% MS-222 (Sigma) and 0.2 mM N-Phenylthiourea (Sigma) and maintained at 28.5°C. The embryos were embedded in a 0.9% agarose column and a 25 −35 μm z stack of the hindbrain, retina, or MHB was acquired with 1 μm steps in a single view, single-sided illumination mode. Images were taken every 0,25 −1 min for 3 - 4 hours using the Plan-Apochromat 40x/1.0 W detection objective (Carl Zeiss Microscopy) and the two PCO.Edge 5.5 sCMOS cameras. The microscope was operated by ZEN 2014 (black edition) software.

### Laser ablations

Nuclei were labelled with H2B-RFP to visualize shifts in chromatin distribution and GFP-PCNA was used as a cell cycle marker. As PCNA is also recruited to sites of DNA damage (Aleksandrov et al., 2018), its enrichment at the ablated region served as a confirmation for successful ablation. Nuclei were ablated in S-phase or several minutes after the onset of G2. A region of interest, consisting of a single point, was selected in the center of each nucleus, resulting in a circular ablated region. 12 μm (16 z-planes) of the tissue surrounding the region of interest were acquired prior to ablation during 3 time points 10 seconds apart. 20 repeats with a frequency of 30 Hz of the laser pulse were performed on the region of interest. 12 μm (16 z-planes) of the tissue surrounding the cut were scanned for 5-15 minutes with temporal resolution of 10 seconds to record the deformations of the ablated region in the hindbrain and the retina after the cut. The shape of the ablated regions was only considered in the first 30 seconds after ablation due to the assumption that later on the damage inflicted by laser ablation might lead to interruption of the force-generating process.

The deformations of the ablated regions were assessed by counting the number of nuclei in which a basal indentation was visible in the z-plane in the center of the nucleus.

### *In situ* hybridization

Riboprobes were generated from cDNA templates and in situ hybridization was performed as previously described (Oates and Ho, 2002; Thisse et al., 1993). The following primers were used to generate the probes for Fmnl3 (lowercase bases contain T7 polymerase promoter):

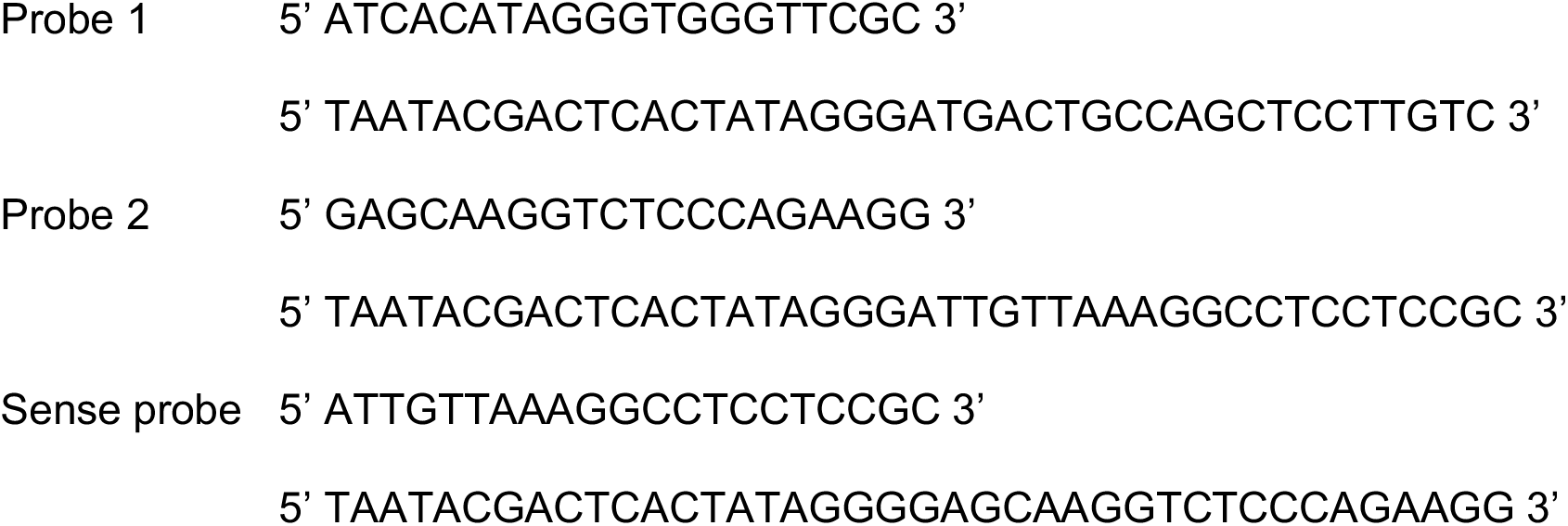

Wholemount stained embryos were documented using a Olympus SZX12 stereomicroscope equipped with a QImaging Micropublisher 5.0 RTV camera (QImaging).

### Image analysis

Minimal image processing was used, prior to image analysis. Processing consisted of image cropping in ZEN and/ or Fiji (Schindelin et al., 2012), bleach correction and/ or background subtraction, as well as drift correction, using Fiji. After image analysis in Imaris 8 or 9 (BitPlane) or Fiji, data was analyzed and plotted using Microsoft Excel, GraphPad Prism or BoxPlotR (Spitzer et al., 2014). Statistical analysis was performed using GraphPad Prism and MATLAB.

#### Sample drift correction

Sample drift in 3D stacks was corrected using a Fiji plugin, created by Benoit Lombardot (Scientific Computing Facility, MPI-CBG, Dresden). The script can be found on http://imagej.net/Manual_drift_correction_plugin.

#### Analysis of apical nuclear migration kinetics

Beginning of G2 was defined by the disappearance of PCNA foci, indicating the end of S-phase, until the onset of cell rounding (Leung et al., 2011). Apical migration was defined by the beginning of directed motion of the nucleus towards the apical side after the onset of G2 and before the onset of cell rounding.

To generate cell trajectories for instantaneous velocities, MSD, and directionality ratio analysis, nuclei were tracked in 3D using Imaris 8 or 9 (Bitplane) during S-phase and G2-phase. Data points were taken at 1 min intervals. The cell axis was defined by the positions of the apical and basal endpoints, measured in 3D in the last time point of S-phase for each cell. Nuclear position was projected onto the cell axis, obtaining one-dimensional time-series, as described (Norden et al., 2009; Leung et al., 2011). Resulting trajectories were analyzed in MATLAB as described previously (Norden et al., 2009; Leung et al., 2011). The kurtosis of instantaneous velocity distributions was calculated using GraphPad Prism.

#### Nuclear segmentation, shape measurements, and tracking in 3D

Semi-automatic segmentation and tracking were performed on 3D stacks in time-series of single labelled migrating nuclei using the Surface tool in Imaris 8 or 9 (Bitplane). The position of the nuclear centroid over time in 3D was extracted. In addition, an ellipsoid was fitted in the segmented surface in each timepoint by the software enabling the extraction of the length of the semi-axes of the nucleus. The nuclear aspect ratio *A/C*) was calculated by dividing the length of each of one of the short semi-axes by the length of the long semi-axis, C.

The average nuclear aspect ratio during S-phase was calculated for each nucleus (*A*/*C*_0_) and used to to calculate the value of the normalised aspect ratio for each timepoint in G2 (*A*/*C*_norm_i__) following the formula:

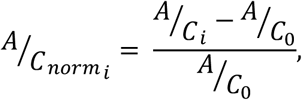

where *A*/*C*_i_ is the nuclear aspect ratio measured in each timepoint (*t*_i_).

Normalized aspect ratios for each time point were pooled for all cells originating from the same tissue.

#### Tissue and cell shape measurements

##### Single cell length measurement in 3D

The 3D viewer of Imaris 8 or 9 (Bitplane) was used to visualize labelled cells in 3D. The positions of the edges of the apical and basal surface were defined using the Measurement Point tool in the last timepoint of S-phase prior to the onset of migration. A custom MATLAB script was used to calculate the distance between the apical and basal surface of each cell and this distance was taken as the length of the cell.

##### Measurement of apical and basal cell footprint areas

Apical and basal cell footprints were segmented semi-automatically in Fiji. Linear regions of interest covering respectively the apical and basal footprint of S-phase cells were drawn to reslice the 3D stack and generate 2D stacks of the cell endfeet of at least 4 timepoints, 1 min apart. The footprints were segmented using automatic thresholding (Huang and Wang, 1995) using the stack histogram, selections were created, and their area measured every minute for 4-10 minutes.

#### Quantifications of actin distribution

##### Tissue-wide actin, myosin, and nuclear distribution profiles

The average fluorescent intensity distribution of phalloidin, phospho-myosin, and DAPI along the apico-basal cell axis was measured as described in (Matejcic et al., 2018; Sidhaye and Norden, 2017). Regions of interest were defined as a 10 μm x 10 μm x (tissue thickness) cuboid for retina and hindbrain and 5 μm x 5 μm x (tissue thickness) cuboid for MHBS and MHBC. 6 to 10 profiles originating from 4-6 samples were measured from each tissue.

##### Actin distribution profiles basally of the nucleus

Basal nuclear actin distribution profiles in hindbrain and retinal cells were measured in Fiji using region of interest immediately basally of the nucleus in each time point. The actin signal intensity in the central z-plane of the cell was measured and normalized to the minimum and the maximum for a given timepoint. The profiles from all timepoints in G2 were used to calculate the average actin profile for the hindbrain and retinal cells.

##### Analysis of basal actin oscillations and cross-correlation analysis

Average basal actin intensity was measured in a square region of interest in the SUM projection of each retinal cell for the duration of G2 using Fiji. Average actin intensity was normalized to the total actin signal in the cell for each timepoint. Data was sampled at intervals of 0.5 min. The rises and plateaus in actin signal, instantaneous velocity, and nuclear aspect ratio were detected and their frequency was calculated as the reciprocal average time difference between the detections, for each timeseries using MATLAB. Both points of increase and stagnation were detected as these were considered to represent biologically relevant maintenance of actin pulses and apical migration. The fluctuations of basal actin were cross-correlated with the fluctuations of mean instantaneous velocity and aspect ratio using MATLAB’s xcorr function. Prior to cross-correlating, each time series was scaled by subtracting its mean and dividing by the standard deviation. A lag of maximal 3 minutes was allowed. Maximum cross-correlation values and the corresponding time lags were used for analysis.

### Statistical analysis

Statistical tests and definitions of error bars are indicated in the figure legends. All statistical tests were two sided. P values > 0.05 were considered not significant. All P values are indicated in the corresponding tables or figure legends. Sample sizes are listed in Tables S3, S4, S5. Statistical analyses were performed using Prism 7 (GraphPad) software and MATLAB.

## Supporting information

Supplemental Material (Model, Figures, Tables)

## ACKNOWLEDGEMENTS

We thank the Norden lab, J. Brugués, S. Grill and especially G. Salbreux for fruitful project discussion. J. Icha, N. Kirkland, Y-L. Mao and P. Strzyz are thanked for helpful comments on the manuscript. We are grateful to H. Hollak, G. Jurado, S. Kaufmann, J. Sidhaye, the Light Microscopy Facility, Scientific Computing and the Fish Facility of the MPI-CBG for experimental help. We thank Li-Kun Phng for sharing the Fmnl3ΔC construct.

I.Y. was a member of the IMPRS of Cell, Developmental and Systems Biology and recipient of an ELBE PhD fellowship. C.N. was supported by MPI-CBG and the German Research Foundation (NO 1068/3-1).

## Author contributions

C.N. and I.Y. conceptualized and decided on methodology used in this work. I.Y. performed majority of experiments and analysis. M.M. helped with experiments. A.E. and M.M. designed analysis tools and performed analysis. C.M. generated the model. C.N. and I.Y. wrote the manuscript with help of all other authors.

