## Supplemental Material (Model, Figures, Tables) for "Cell and tissue morphology determine actin-dependent nuclear migration mechanisms in neuroepithelia"

### Proof-of-Principle Model of a Nuclear Migration Mechanism in Retinal Neuroepithelia

#### 1 Critical Buckling & Saltatory Dynamics

We first need to calculate the critical buckling length for an f-actin filament or bundle. We use as many measured or approximate community-accepted values as possible and can "fit" or extract information about those that remain free from the observed dynamics. We also here assume that the timescale for any viscoelastic relaxation of the f-actin bundles is slower than the relevant buckling dynamics. For classical Euler buckling, the critical buckling length is given by [1]:

$$L_c = c_{bc}\pi \left(\frac{\kappa_B}{F}\right)^{1/2} \tag{1}$$

for $c_{bc} = 1/2$ in the case of one clamped and one free end of the filament. $\kappa_B$ is the bending stiffness of the filament or bundle and $F$ is the applied compressional force. For the applied force, we are deep into the regime of low Reynolds number and may assume a Stokes' drag scenario for the nucleus being pushed through the cytoplasm by the f-actin. This gives the following equation for $F$:

$$F = 6\pi\eta R v \tag{2}$$

for $\eta$ the effective dynamic viscosity of cytoplasm at the length scale of the nucleus, $R$ the radius of the nucleus, and $v$ the flow velocity. Meanwhile, for bundled f-actin filaments the bending stiffness is given by [2]:

$$\kappa_B = \kappa_f N \left[1 + \frac{A_f(N-1)(d_f+t)^2/12I_f}{1 + c(q_j)\frac{N+\sqrt{N}}{\alpha}}\right]. \tag{3}$$

Here $\kappa_f$ is the bending stiffness of a single strand of f-actin, $N$ is the number of filaments in the bundle, and $A_f, d_f, t,$ and $I_f$ are geometric properties of the actin monomers and the cross-linker spacings. $c(q_j) \approx 1$ for the low relevant mode numbers with our boundary conditions. $\alpha$ is a unitless parameter that measures the relative importance of the cross-linker shear stiffness, $\alpha = 0$ in the limit of low shear stiffness and $\alpha \to \infty$ for very high shear stiffness. If we assume that myosin is coordinating and cross-linking the bundles, then $\alpha$ is likely to be very low and we may approximate $\kappa_B \approx \kappa_f N$.

To show a proof-of-principle for the hypothesis that the buckling is relevant for the migration mechanics, we would expect a plausible set of these parameters in the above equations to allow for an $L_c$ of order $1-10\mu m$, which would match the experimentally observed distances between the anchoring formin domain and the basal side of the nucleus. The following values were approximated for the needed parameters:

$$\eta \approx 5 * 10^{-2}\, Pa\, sec$$
$$R \approx 3 * 10^{-6}\, m$$
$$v \approx 1.6 * 10^{-8}\, m/sec$$
$$\kappa_f \approx 10^{-25}\, Nm^2$$

with $\eta$ and $\kappa$ taken from the literature [3, 4] and $R$ and $v$ taken from our own observations.

Under these approximations we indeed find that $L_c$ is plausibly in the neighborhood of $1 - 10\mu m$, consistent with the observations regarding the trailing formin attachments. $N$ can easily be taken to be up to 20 or so without pushing $L_c$ out of a plausible range. Furthermore, increasing the shear stiffness of the cross-linkers in any putative bundling would only serve to push the overall bending stiffness of the bundle out of range on the top end, providing some circumstantial evidence that the shear-soft myosins serve as the primary cross-linkers.

The sensitivity of $L_c$ to the effective dynamic viscosity allows our proof-of-principle model to qualitatively explain the saltatory motion of the nucleus as well. In the crowded environment of the PSE, the effective $\eta$ seen by the traveling nucleus can vary widely as intra-cellular machinery or compartments are encountered or stiffer, interposing regions from neighboring cells, associated, for example, with the locations of their nuclei crowd the local environment. All of these things can serve to temporarily increase the compressive force acting on the actin filaments and bundles, leading to lower critical buckling threshold lengths. As the filaments buckle, continued polymerization at the formin-anchored end no longer moves the nucleus forward but instead increases the curvature of the buckled filament, adding to the stored stress and hence the force delivered to the nucleus. Eventually, the rising stress in the buckled filaments provides enough force to push the nucleus past whatever local road block caused the buckling in the first place. When this occurs, the filaments can then rapidly straighten, causing a burst of increased velocity for the nucleus. It is likely that the force required to resolve these impediments is directly related to the deformability of the nuclear envelope, potentially explaining why Lamin A over expression slowed migration, as greater forces on the nucleus and stresses in the filaments would be required, increasing the frequency with which individual filaments might undergo critical failure and subsequent depolymerization.

This picture of f-actin growth pushing the nucleus forward with alternating periods in which the polymerization leads directly to increased displacement of the nucleus and in which it leads instead to rising forces delivered to a stalled nucleus does make one key prediction. Because the motion of the nucleus is driven directly by the polymerization, and the displacement after stalls are resolved is again simply the length of the actin bundle, then the average speed of the nucleus over its migratory period should very closely match the speed of f-actin polymerization. And indeed, formin-catalyzed f-actin polymerization is known to be approximately [5]:

$$k_+ \approx 0.3\, \mu m/min.$$

which is very much in line with our observations of the average velocity of the migrating nucleus.

### Supplementary Figure Legends

**Figure S1. Hindbrain and retinal nuclei move and deform differently during apical nuclear migration.** Related to Fig. 1 and 2. (**A**) Mean squared displacement of hindbrain and retinal cell apical migration. Mean of all tracks is shown, error bars: SD. (**B**) Control staining of Lamin A/C in the tail of a 24 hpf zebrafish (lookup tables indicate minimal and maximal Lamin B1 and Lamin A/C signal values). Scale bar: 10 µm. (**C**) Normalized aspect ratio changes of individual hindbrain and retinal nuclei with time. (**D**) Fluctuations of mean instantaneous velocity and normalized nuclear aspect ratio with time in representative retinal cell. (**E**) Schematic of the nuclear laser ablation experiments. A circular region is ablated in the center of hindbrain and retinal nuclei. Changes in the shape of the ablated region are analyzed to infer a pulling or pushing force acting on nuclei. (**F**) Representative image of ablated neuroepithelial nucleus. H2B-RFP (gray) labels chromatin and GFP-PCNA (green) serves as a cell cycle phase marker. The shape of the ablated region is assessed in the H2B channel while successful ablation can be verified in the PCNA channel by the accumulation of PCNA in regions of DNA damage. (**G**) Representative images of hindbrain and retinal nuclei in S or G2 phase immediately after laser ablation. (**H**) Summary table of the effects of nuclear laser ablation on nuclei in different tissues and different cell cycle phases. Shortening and basal indentation are only observed in ablated regions of retinal G2 nuclei.

**Figure S2. Apical nuclear migration in the hindbrain depends on actomyosin but myosin distribution differs in hindbrain and retinal cells.** Related to Fig. 3. (**A**)-(**D**) Representative time-series of cells treated with different cytoskeletal inhibitors in hindbrain and retina (Video 3). Samples were incubated in DMSO, 100 µM Colcemid (microtubule polymerization inhibitor), 100 µM Blebbistatin (myosin II inhibitor), or 2 μM Latrunculin A (actin polymerization inhibitor) and 8 μM Jasplakinolide (microfilament turnover inhibitor). (**E**) Normalized average intensity distribution of GFP-UtrophinCH signal. Shown mean profile of all S-phase time points, error bars: SD. (**F**) Myosin distribution upon apical migration in a single hindbrain (upper) and retinal (lower) cell. mKate2-PCNA labels nuclei (gray), CA-MRLC-GFP labels active myosin (lookup table indicates minimal and maximal CA-MRLC-GFP signal values). (**G**) Normalized average intensity distribution of CA-MRLC-GFP signal. Shown mean profile of all G2 time points, error bars: SD. (**H'**, **H''**) Fluctuations in normalized basal actin plotted together with fluctuations in instantaneous velocity (**H'**) and normalized nuclear aspect ratio (**H''**) in same retinal cell. (**I**) Pooled frequencies of oscillation of instantaneous velocity, nuclear aspect ratio, and basal actin intensity for seven retinal cells. White circles show the means; box limits indicate the 25th and 75th percentiles as determined by R software; whiskers extend 1.5 times the interquartile range from the 25th and 75th percentiles; polygons represent density estimates of data and extend to extreme values. Scale bars: 5 μm.

**Figure S3. Apical nuclear migration in hindbrain and retina is controlled by different actomyosin regulators.** Related to Fig. 4. (**A**)-(**C**) Representative time-series of cells treated with different inhibitors of actomyosin dynamics in hindbrain and retina (Video 6). Samples were incubated in (**A**) 200 μM Rhosin and 50 μM Y16 (RhoA-GTPase inhibitors), (**B**) 250 μM ML-7 (MLCK inhibitor), or (**C**) 200 μM CK-666 (Arp2/3 inhibitor). Scale bars: 5 μm. (**D**) Table summarizing the effects of actomyosin dynamics perturbation experiments on apical nuclear migration in hindbrain and retina.

**Figure S4. Actin polymerization factors in apical migration and links between tissue morphology and actin-dependent force generation.** Related to Fig. 4, 5, and 6. (**A**), (**B**) Representative time-series of hindbrain and retinal cells expressing (**A**) control mKate2-Ras and GFP-PCNA (both in gray) or (**B**) heat-shock induced dominant negative mKate2-N-WASP-CA and GFP-PCNA (both in gray) (Video 7). N-WASP inhibition did not perturb apical migration in hindbrain. (**C**) *In situ* hybridization analysis demonstrates that *fmnl3* is expressed in the retina at 24 hpf. (**D**) Cell length of hindbrain, retinal, MHBS, and MHBC cells (Table 2). $p_{(HB\text{-}MHBS)}=0.1017$ , $p_{(R\text{-}MHBC)}=0.0053$ , $p_{(HB\text{-}R)}=0.5780$ , $p_{(MHBS\text{-}MHBC)}=0.0872$. (**E**) Apical-to-basal footprint ratio of hindbrain, retinal, MHBS, and MHBC cells (Table 2). $p_{(HB\text{-}MHBS)}=0.1312$ , $p_{(R\text{-}MHBC)}=0.2617$ , $p_{(HB\text{-}R)}<0.0001$ , $p_{(MHBS\text{-}MHBC)}<0.0001$. (**F**) Duration of apical migration of hindbrain, retinal, MHBS, and MHBC cells (Table 2). $p_{(HB\text{-}MHBS)}=0.3540$ , $p_{(R\text{-}MHBC)}=0.2457$ , $p_{(HB\text{-}R)}= 0.0052$ , $p_{(MHBS\text{-}MHBC)}=0.0326$. (**G**) Instantaneous velocity distribution of hindbrain, retinal, MHBS, and MHBC cells (Table 2). $p_{(HB\text{-}MHBS)}=0.9423$ , $p_{(R\text{-}MHBC)}=0.7831$ , $p_{(HB\text{-}R)}<0.0001$ , $p_{(MHBS\text{-}MHBC)}=0.0038$. All error bars in (**D**)-(**G**): SD. All p values in (**D**)-(**G**) calculated using Mann-Whitney test. (**H**) Representative time-series of MHBS and MHBC cells treated with different 125 µM Rockout (ROCK inhibitor). Scale bars**:** 5 µm.

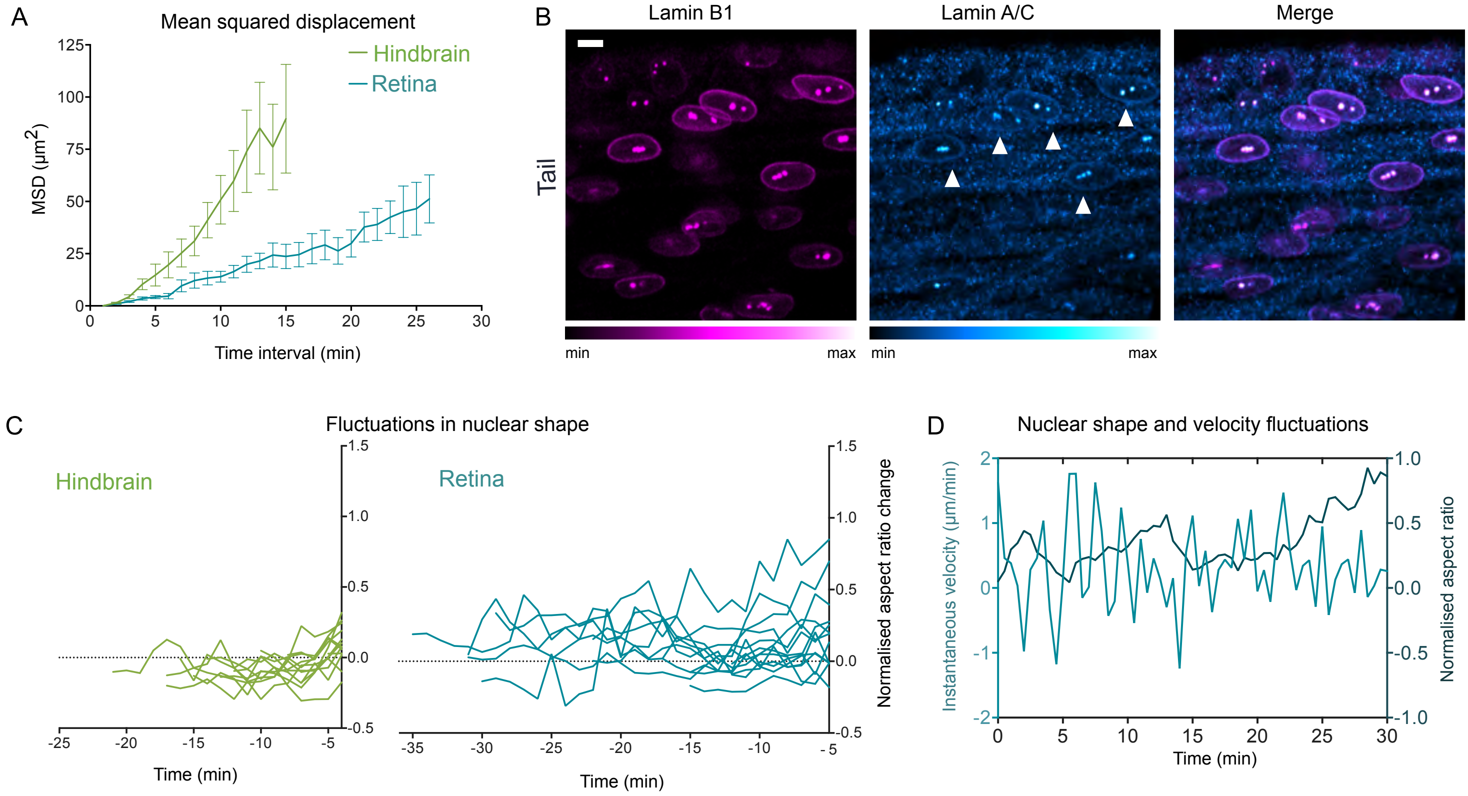


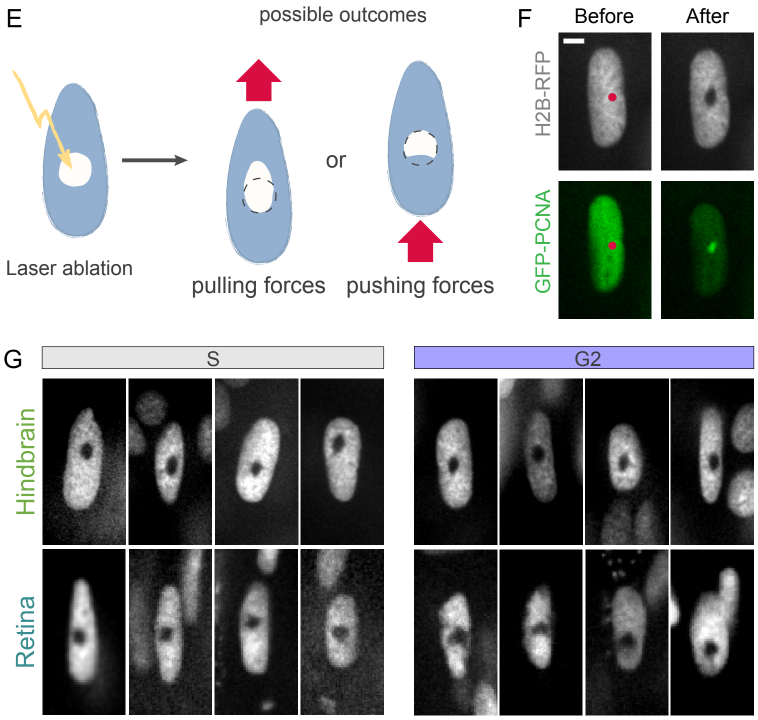


H

| | Hindbrain (S) | Hindbrain (G2) | Retina (S) | Retina (G2) |
|---|---|---|---|---|
| Deformation events | 0 out of 12 | 0 out of 10 | 0 out of 10 | 7 out of 14 |

**Figure S1. Hindbrain and retinal nuclei move and deform differently during apical nuclear migration.** Related to Fig. 1 and 2. (**A**) Mean squared displacement of hindbrain and retinal cell apical migration. Mean of all tracks is shown, error bars: SD. (**B**) Control staining of Lamin A/C in the tail of a 24 hpf zebrafish (lookup tables indicate minimal and maximal Lamin B1 and Lamin A/C signal values). Scale bar: 10 μm. (**C**) Normalized aspect ratio changes of individual hindbrain and retinal nuclei with time. (**D**) Fluctuations of mean instantaneous velocity and normalized nuclear aspect ratio with time in representative retinal cell. (**E**) Schematic of the nuclear laser ablation experiments. A circular region is ablated in the center of hindbrain and retinal nuclei. Changes in the shape of the ablated region are analyzed to infer a pulling or pushing force acting on nuclei. (**F**) Representative image of ablated neuroepithelial nucleus. H2B-RFP (gray) labels chromatin and GFP-PCNA (green) serves as a cell cycle phase marker. The shape of the ablated region is assessed in the H2B channel while successful ablation can be verified in the PCNA channel by the accumulation of PCNA in regions of DNA damage. (**G**) Representative images of hindbrain and retinal nuclei in S or G2 phase immediately after laser ablation. (**H**) Summary table of the effects of nuclear laser ablation on nuclei in different tissues and different cell cycle phases. Shortening and basal indentation are only observed in ablated regions of retinal G2 nuclei.

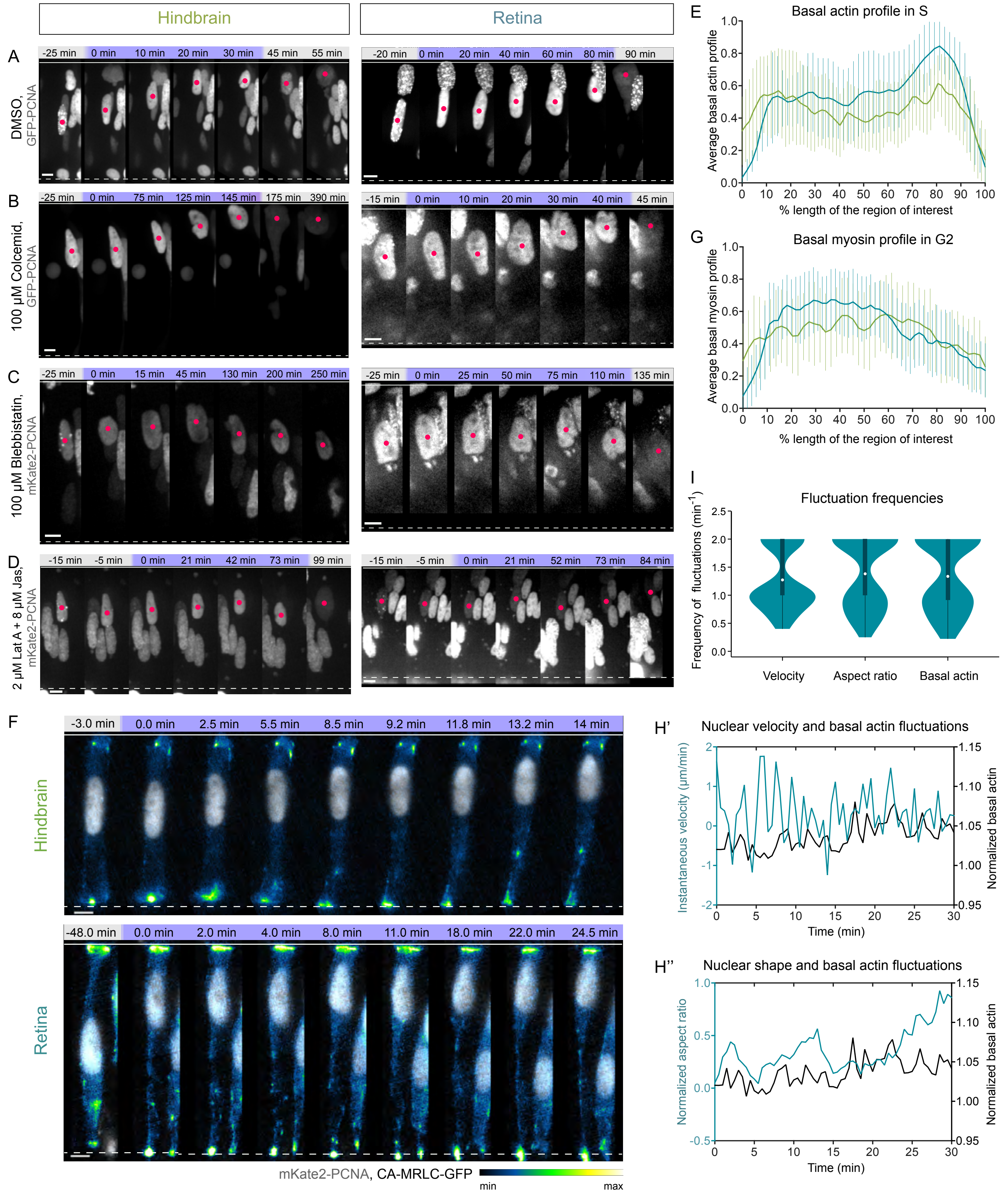


**Figure S2. Apical nuclear migration in the hindbrain depends on actomyosin but myosin distribution differs in hindbrain and retinal cells.** Related to Fig. 3. (**A**)-(**D**) Representative time-series of cells treated with different cytoskeletal inhibitors in hindbrain and retina (Video 3). Samples were incubated in DMSO, 100 µM Colcemid (microtubule polymerization inhibitor), 100 µM Blebbistatin (myosin II inhibitor), or 2 µM Latrunculin A (actin polymerization inhibitor) and 8 µM Jasplakinolide (microfilament turnover inhibitor). (**E**) Normalized average intensity distribution of GFP-UtrophinCH signal. Shown mean profile of all S-phase time points, error bars: SD. (**F**) Myosin distribution upon apical migration in a single hindbrain (upper) and retinal (lower) cell. mKate2-PCNA labels nuclei (gray), CA-MRLC-GFP labels active myosin (lookup table indicates minimal and maximal CA-MRLC-GFP signal values). (**G**) Normalized average intensity distribution of CA-MRLC-GFP signal. Shown mean profile of all G2 time points, error bars: SD. (**H'**, **H''**) Fluctuations in normalized basal actin plotted together with fluctuations in instantaneous velocity (**H'**) and normalized nuclear aspect ratio (**H''**) in same retinal cell. (**I**) Pooled frequencies of oscillation of instantaneous velocity, nuclear aspect ratio, and basal actin intensity for seven retinal cells. White circles show the means; box limits indicate the 25th and 75th percentiles as determined by R software; whiskers extend 1.5 times the interquartile range from the 25th and 75th percentiles; polygons represent density estimates of data and extend to extreme values. Scale bars: 5 µm.

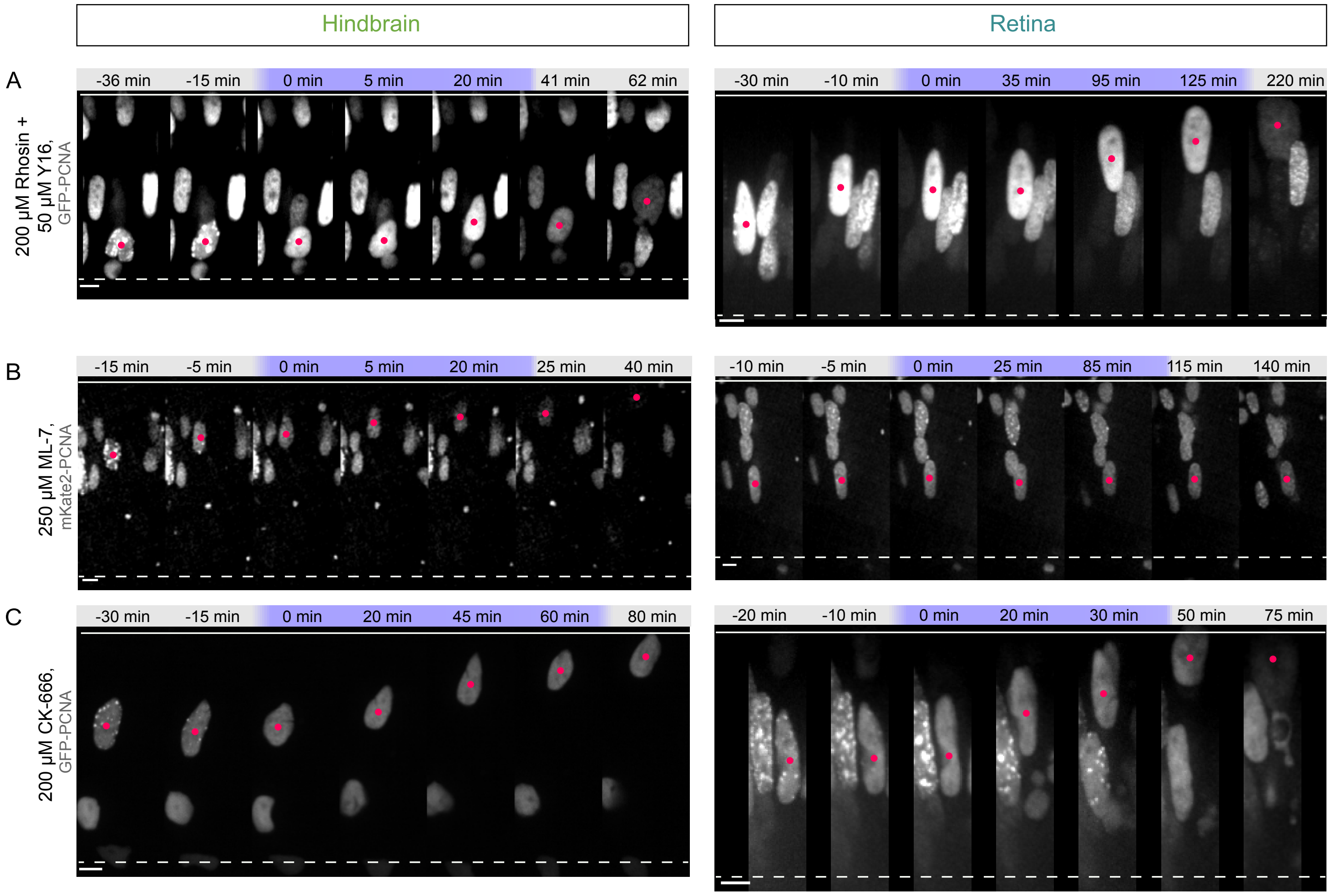


D

| Small molecule inhibitor | Inhibited process | Apical migration perturbed | |
|---|---|---|---|
| | | Hindbrain | Retina |
| Colcemid | Microtubule polymerization | no | no |
| Blebbistatin | Myosin II activity | yes | yes |
| Jasplakinolide + Latrunculin A | Actin filament turnover | yes | yes |
| Rhosin + Y16 | Rho-GTPase activity | yes | no |
| Rockout | Rho-kinase activity | yes | no |
| ML-7 | MLCK activity | no | yes |
| CK-666 | Arp2/3-dependent actin nucleation | no | no |
| SMIFH2 | Formin-dependent actin nucleation | no | yes |

**Figure S3. Apical nuclear migration in hindbrain and retina is controlled by different actomyosin regulators.** Related to Fig. 4. (**A**)-(**C**) Representative time-series of cells treated with different inhibitors of actomyosin dynamics in hindbrain and retina (Video 6). Samples were incubated in (**A**) 200 µM Rhosin and 50 µM Y16 (RhoA-GTPase inhibitors), (**B**) 250 µM ML-7 (MLCK inhibitor), or (**C**) 200 µM CK-666 (Arp2/3 inhibitor). Scale bars: 5 µm. (**D**) Table summarizing the effects of actomyosin dynamics perturbation experiments on apical nuclear migration in hindbrain and retina.

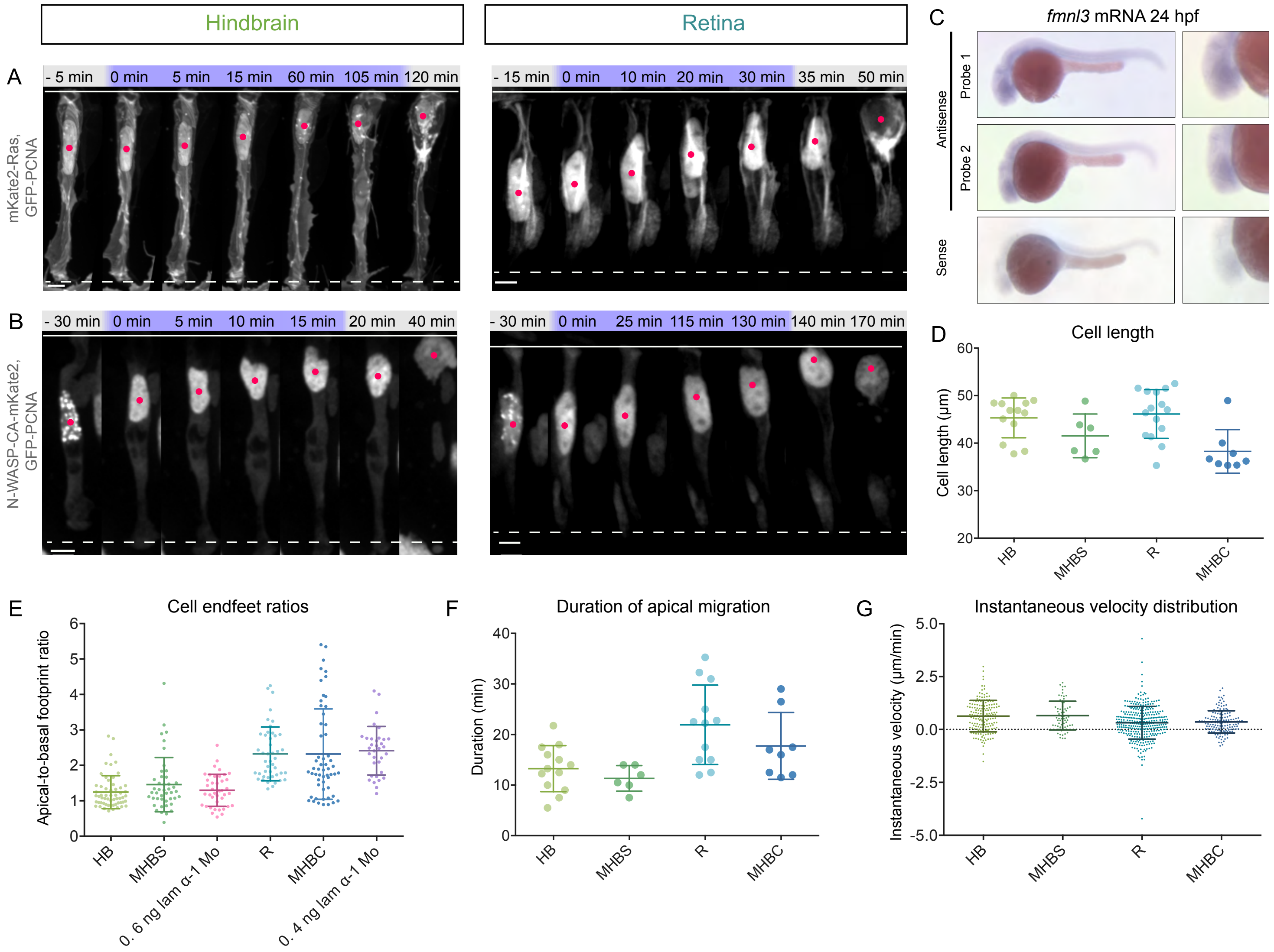


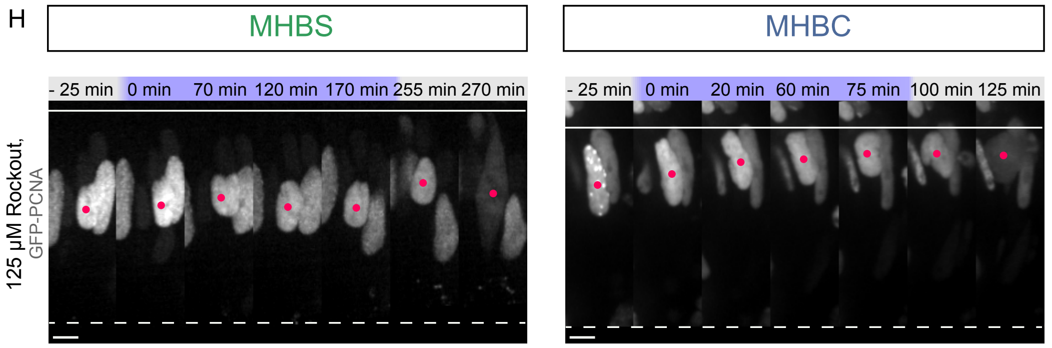


**Figure S4. Actin polymerization factors in apical migration and links between tissue morphology and actin-dependent force generation.** Related to Fig. 4, 5, and 6. (**A**), (**B**) Representative time-series of hindbrain and retinal cells expressing (**A**) control mKate2-Ras and GFP-PCNA (both in gray) or (**B**) heat-shock induced dominant negative mKate2-N-WASP-CA and GFP-PCNA (both in gray) (Video 7). N-WASP inhibition did not perturb apical migration in hindbrain. (**C**) *In situ* hybridization analysis demonstrates that *fmnl3* is expressed in the retina at 24 hpf. (**D**) Cell length of hindbrain, retinal, MHBS, and MHBC cells (Table 2). $p_{(HB\text{-}MHBS)}=0.1017$ , $p_{(R\text{-}MHBC)}=0.0053$ , $p_{(HB\text{-}R)}=0.5780$ , $p_{(MHBS\text{-}MHBC)}=0.0872$. (**E**) Apical-to-basal footprint ratio of hindbrain, retinal, MHBS, and MHBC cells (Table 2). $p_{(HB\text{-}MHBS)}=0.1312$ , $p_{(R\text{-}MHBC)}=0.2617$ , $p_{(HB\text{-}R)}<0.0001$ , $p_{(MHBS\text{-}MHBC)}<0.0001$. (**F**) Duration of apical migration of hindbrain, retinal, MHBS, and MHBC cells (Table 2). $p_{(HB\text{-}MHBS)}=0.3540$ , $p_{(R\text{-}MHBC)}=0.2457$, $p_{(HB\text{-}R)}= 0.0052$ , $p_{(MHBS\text{-}MHBC)}=0.0326$. (**G**) Instantaneous velocity distribution of hindbrain, retinal, MHBS, and MHBC cells (Table 2). $p_{(HB\text{-}MHBS)}=0.9423$ , $p_{(R\text{-}MHBC)}=0.7831$ , $p_{(HB\text{-}R)}<0.0001$ , $p_{(MHBS\text{-}MHBC)}=0.0038$. All error bars in (**D**)-(**G**): SD. All p values in (**D**)-(**G**) calculated using Mann-Whitney test. (**H**) Representative time-series of MHBS and MHBC cells treated with different 125 μM Rockout (ROCK inhibitor). Scale bars: 5 μm.

Supplementary tables to

### Cell and tissue morphology determine actin-dependent nuclear migration mechanisms in neuroepithelia

Iskra Yanakieva, Anna Erzberger, Marija Matejčić, Carl D. Modes and Caren Norden

| Construct | Labelled structure/ Function | Reference |
|---|---|---|
| hsp70: Ras-mKate2 | Cell membrane | (Strzyz et al., 2015) |
| bactin: mKate2-Ras | Cell membrane | (Icha et al., 2016) |
| hsp70: GFP-UtrophinCH | F-actin | (Strzyz et al., 2015) |
| hsp70: PCNA-GFP | Cell cycle phase marker | (Icha et al., 2016) |
| hsp70: DN-Rok2-EGFP/ mKate | Dominant negative Rho-kinase | This study |
| hsp70: mKate2-N-WASP-CA | C-terminal domain of the Arp2/3 activator N-WASP | (Icha et al., 2016) |
| hsp70: EGFP-LAP2b | Lamina-associated polypeptide 2 | This study |
| hsp70: LMNA-mKate2 | Lamin A | This study |
| hsp70: Fmnl3-EGFP | Fmnl3 | This study |
| hsp70: Fmnl3ΔC-EGFP | Truncated Fmnl3 | Recloned from (Phng et al., 2015) |
| pCS2+ Ras-mKate2 | Cell membrane | (Weber et al., 2014) |
| pCS2+ Ras-GFP | Cell membrane | (Strzyz et al., 2015) |
| pCS2+ H2B-RFP | Chromatin | (Norden et al., 2009) |
| pCS2+ GFP-UtrophinCH | F-actin | (Burkel et al., 2007) |
| pCS2+ Lifeact-GFP | F-actin | Kind gift from Oates lab |
| pCS2+ MRLC2$^{T18DS19D}$-GFP | Constitutively activated myosin light chain | (Strzyz et al., 2015) |
| pCS2+ PCNA-GFP | Cell cycle phase marker | (Leung, Klopper et al., 2011) |
| pCS2+ mKate2-PCNA | Cell cycle phase marker | This study |

**Supplementary Table 1.** List of constructs.

| Chemical | Inhibited process | Working concentration | Source/ Cat.No. |
|---|---|---|---|
| Blebbistatin | Myosin II activity | 100 µM | Enzo Life Sciences/ BML-EI315-0005 |
| Colcemid | Microtubule polymerization | 100 µM | Enzo Life Sciences/ ALX-430-033-M005 |
| Jasplakinolide | Actin filament turnover | 8 µM | BIOMOL Feinchemikalien/ AG-CN2-0037-C050 |
| Latrunculin A | Actin filament polymerization | 2 µM | GmbH/ 10010630 |
| Rhosin | Rho-GTPase activity | 200 µM | Merck Millipore/ 555460 |
| Y16 | Rho-GTPase activity | 50 µM | Sigma Aldrich/ SML0873 |
| Rockout | Rho-kinase activity | 125 µM | Santa Cruz Biotechnology/ sc-203237 |
| ML-7 | MLCK activity | 250 µM | Enzo Life Sciences/ BML-EI197-0010 |
| CK-666 | Arp2/3-dependent actin nucleation | 200 µM | Merck Millipore/ 182515 |
| SMIFH2 | Formin-dependent actin nucleation | 10 µM | Merck Millipore/ 344092 |

**Supplementary Table 2.** List of cytoskeleton inhibitors.

| Experiment | Number of embryos (N) | | Number of cells (n) | | Statistical test | Figure |
|---|---|---|---|---|---|---|
| | Hindbrain | Retina | Hindbrain | Retina | | |
| MSD | 5 | 10 | 13 | 15 | - | Suppl. 1a |
| Migration starting points (variance) | 5 | 10 | 13 | 15 | F test | 1e |
| Nuclear velocity – aspect ratio cross-correlation | 4 | 10 | 13 | 13 | Wilcoxon signed-rank | 2h |
| Fluctuation frequencies (velocity, aspect ratio, basal actin) | - | 3 | - | 4 | Mann Whitney | Suppl. 2c |
| Nuclear velocity, aspect ratio, basal actin cross-correlation | - | 3 | - | 4 | - | 3e |
| DMSO control (Rockout) | 6 | 5 | 131 | 105 | Mann Whitney | 4a, d |
| Rockout live treatment | 6 | 10 | 250 | 141 | Mann Whitney | 4b, d |
| DMSO control (SMIFH2) | 4 | 8 | 18 | 61 | Mann Whitney | 4e |
| SMIFH2 live treatment | 4 | 10 | 23 | 57 | Mann Whitney | 4c, e |
| Unaffected Rockout live treatment | 2 | - | - | - | - | - |
| Unaffected SMIFH2 live treatment | - | 16 | - | - | - | - |

**Supplementary Table 3.** Number of embryos and cells used in the analysis comparing different parameters in hindbrain and retina.

| Experiment | Number of embryos (N) | | | | Number of cells (n) | | | | Statistical test | Figure |
|---|---|---|---|---|---|---|---|---|---|---|
| | HB | R | MHBS | MHBC | HB | R | MHBS | MHBC | | |
| Cell length | 5 | 10 | 5 | 7 | 13 | 15 | 6 | 8 | Mann Whitney | Table 2 |
| Apical-to-basal footprint ratio | 4 | 4 | 4 | 7 | 6 | 5 | 5 | 7 | Mann Whitney | Table 2 |
| Duration of apical migration (min) | 5 | 10 | 4 | 7 | 13 | 15 | 5 | 8 | Mann Whitney | Table 1, 2 |
| Mean instantaneous velocity (µm/min) | 5 | 10 | 4 | 7 | 13 | 15 | 5 | 8 | Mann Whitney | Table 1, 2 |
| Basal-to-apical distribution profiles | 7 | 6 | 10 | 8 | - | - | - | - | - | 5a, b, d, e |
| Nuclear trajectories | 5 | 10 | 4 | 7 | 13 | 15 | 5 | 8 | - | 1c, d, 5f |
| Directionality ratio | 5 | 10 | 4 | 7 | 13 | 15 | 5 | 8 | - | 1f, 5g |
| Normalized nuclear aspect ratio | 4 | 10 | 5 | 5 | 13 | 13 | 6 | 6 | Mann Whitney | 2g, Suppl. 1c, 5h |

**Supplementary Table 4.** Number of embryos and cells used in the analysis comparing different parameters in hindbrain, retina, MHBS, and MHBC.

| Representative data | Number of instances observed | | Figure |
|---|---|---|---|
| | Hindbrain | Retina | |
| Lamin B1, Lamin A/C staining | 3 | 3 | 2a |
| Nuclear deformations (LAP2b) | - | 3 | 2b |
| Nuclear deformations upon laser ablation | 1 | 1 | 2c |
| Perturbed apical migration upon LmnA overexpression | - | 3 | 2e |
| Actin distribution in single G2 cells | 3 | 4 | 3a |
| Basal actin profile | 1 | 1 | 3b’ |
| Basal actin fluctuations | - | 3 | 3c’ |
| DN-Rok2-EGFP | 5 | 10 | 3f |
| Fmnl3ΔC-EGFP | 14 | 19 | 3g |
| Fmnl3-EGFP distribution | - | 3 | 3h |
| Myosin distribution in single G2 cells | 5 | 3 | Suppl. 2b |
| N-WASP-CA-mKate2 | 15 | 11 | Suppl. 4b |
| Fmnl3 in situ hybridization | - | 2 independent experiments | Suppl. 4c |

**Supplementary Table 5.** Number of instances different unquantified observations were made.
